# SARS-CoV-2 Nsp14 binds Tollip and activates pro-inflammatory pathways while downregulating interferon-α and interferon-γ receptors

**DOI:** 10.1101/2024.12.12.628214

**Authors:** Naveen Thakur, Poushali Chakraborty, JoAnn M. Tufariello, Christopher F. Basler

**Author notes:** Corresponding authors: Christopher F. Basler, Department of Microbiology, Icahn School of Medicine at Mount Sinai, New York, NY 10029,; JoAnn M. Tufariello, Department of Microbiology, Icahn School of Medicine at Mount Sinai, New York, NY 10029.

## Abstract

SARS coronavirus 2 (SARS-CoV-2) non-structural protein 14 (Nsp14) possesses an N-terminal exonuclease (ExoN) domain that provides a proofreading function for the viral RNA-dependent RNA polymerase and a C-terminal N7-methyltransferase (N7-MTase) domain that methylates viral mRNA caps. Nsp14 also modulates host functions. This includes the activation of NF-κB and downregulation of interferon alpha/beta receptor 1 (IFNAR1). Here we demonstrate that Nsp14 exerts broader effects, activating not only NF-κB responses but also ERK, p38 and JNK MAP kinase (MAPK) signaling, promoting cytokine production. Further, Nsp14 downregulates not only IFNAR1 but also IFN-γ receptor 1 (IFNGR1), impairing cellular responses to both IFNα and IFNγ. IFNAR1 and IFNGR1 downregulation is via a lysosomal pathway and also occurs in SARS-CoV-2 infected cells. Analysis of a panel of Nsp14 mutants reveals a consistent pattern. Mutants that disable ExoN function remain active, whereas N7-MTase mutations impair both pro-inflammatory pathway activation and IFN receptor downregulation. Innate immune modulating functions also require the presence of both the ExoN and N7-MTase domains likely reflecting the need for the ExoN domain for N7-MTase activity. We further identify multi-functional host protein Tollip as an Nsp14 interactor. Interaction requires the phosphoinositide-binding C2 domain of Tollip and sequences C-terminal to the C2 domain. Full length Tollip or regions encompassing the Nsp14 interaction domain are sufficient to counteract both Nsp14-mediated and Nsp14-independent activation of NF-κB. Knockdown of Tollip partially reverses IFNAR1 and IFNGR1 downregulation in SARS-CoV-2 infected cells, suggesting relevance of Nsp14-Tollip interaction for Nsp14 innate immune evasion functions.

**Significance:** SARS-CoV-2 protein Nsp14 both activates NF-κB, which promotes virus replication and inflammation, and downregulates IFNAR1, which can render infected cells resistant to the antiviral effects of IFN-α/β. Our study demonstrates that Nsp14 also activates MAPK signaling and downregulates IFNGR1, causing broader impacts than previously recognized. Data from a panel of Nsp14 mutants suggests a common underlying effect of Nsp14 may be responsible for its multiple innate immune activities. We further describe a novel interaction between Nsp14 and Tollip, a selective autophagy receptor. We show that Tollip expression downregulates Nsp14 activation of NF-κB and that Tollip knockdown reverses IFNAR1 and IFNGR1 downregulation in SARS-CoV-2 infection, suggesting that Tollip functions as a regulator of Nsp14 innate immune modulation.

## Introduction

Severe acute respiratory syndrome coronavirus 2 (SARS-CoV-2) is the cause of the coronavirus disease 2019 (COVID-19) pandemic which has been responsible for over 776 million reported cases and 7.06 million deaths worldwide since 2019 (1). SARS-CoV-2 is an enveloped virus with a positive-sense, single-stranded RNA genome of ∼30 kb (2). The genome encodes 16 non-structural proteins, the structural proteins spike (S), envelope (E), membrane (M), and nucleocapsid (N), and several accessory proteins (3). Among the non-structural proteins is Nsp14, a bifunctional protein with well-established roles in viral RNA synthesis. Nsp14 possesses two distinct domains, one at the N-terminus with exoribonuclease (ExoN) activity that removes nucleotide misincorporations during viral RNA synthesis (4). Nsp14 exonuclease activity is greatly enhanced by the addition of Nsp10, which interacts with the N-terminus of Nsp14, promoting conformational change and enhancing stability (5–7). Due to the high error rate of the coronavirus RNA-dependent-RNA-polymerase (RdRp) (8), the proofreading activity provided by the Nsp10-Nsp14 complex is critical for maintaining the integrity of the large genome (4, 9–11). The second functional domain, found at the Nsp14 C-terminus, is a guanine-N7-methyltransferase (N7-MTase) which contributes to viral RNA cap methylation and to virus replication (12–14). This capping promotes translation of the coronavirus mRNA and interferes with innate immune recognition (15, 16).

SARS-CoV-2 modulates innate immunity in a variety of ways that promote viral infection and disease. For example, SARS-CoV-2 infection activates NF-κB and mitogen activated protein kinase (MAPK) signaling (17–19). Activation of NF-κB is needed for SARS-CoV-2 infection as indicated by reduced viral protein production and infection when NF-κB signaling was inhibited by knockdown or small molecule inhibitor approaches (17). Inhibition of p38 MAPK impairs replication, demonstrating a pro-SARS-CoV-2 role for the pathway (18, 19). Knockdown of MAP2K4/MKK4, which activates JNK, also inhibited viral replication suggesting a role for this arm of MAPK signaling in SARS-CoV-2 infection (19). Activation of these pathways may contribute to uncontrolled inflammatory responses and cytokine storm that in some cases results in acute respiratory distress syndrome (ARDS), multiorgan failure, and fatal outcomes among COVID-19 patients (20–22). SARS-CoV-2 also antagonizes IFN responses, with a wide variety of mechanisms reported (23). Some of the mechanisms are due to general effects on host cell processes. For example, coronaviruses shut down host protein synthesis through the actions of multiple viral proteins (24–28). In particular, Nsp1 blocks translation by promoting cellular mRNA degradation and by binding to the 40S ribosomal subunit; and it blocks mRNA nuclear export (25, 26, 29–35). ORF6 interacts with importin alpha and nucleoporins, disrupting nuclear-cytoplasmic trafficking (36–42). These activities suppress IFN responses. Other viral proteins may target specific innate immune pathways (23).

Besides its well-characterized roles in proofreading and mRNA cap methylation, Nsp14 has also been implicated in modulation of innate immunity. Overexpression of Nsp14 causes near total shutdown of cellular protein synthesis, including synthesis of antiviral IFN-stimulated genes (ISGs) (43). This effect is reversed by mutations that inactivate either the ExoN or the N7-MTase activity and is enhanced by Nsp10 (43). Nsp14 has also been linked to global dysregulation of host transcription and inhibition of host mRNA processing and mRNA nuclear export (44, 45), activities that likely modulate host innate immune responses to infection. Nsp14 has also been implicated in the activation of NF-κB and the suppression of IFNα/β signaling. Nsp14 activation of NF-κB has been attributed to Nsp14 interaction with inosine-5’-monophosphate dehydrogenase 2 (IMPDH2); through interaction with the IKK complex (IKKα/IKKβ/NEMO), which may be recruited due to linear ubiquitination of Nsp14, or by a mechanism that does not involve interaction with the IKK complex (IKKα/IKKβ/NEMO) (45–51). Mutations designed to abrogate MTase activity impair NF-κB activation, but in one study N7-MTase inhibitors (nitazoxanide and sinefungin) did not block NF-κB signaling (47). More recently, Nsp14 was described to promote MAP kinase signaling, specifically activating ERK and triggering of AP-1-dependent gene expression (52). Beyond activation of innate immune pathways, screens to identify SARS-CoV-2 interferon antagonists identified Nsp14 as an inhibitor of IFNβ promoter induction and IRF3 nuclear accumulation (53, 54). Nsp14 antagonizes IFNα/β induced signaling by downregulating expression of IFNAR1, one of two proteins that comprise the IFNα/β receptor, possibly through lysosomal degradation (36, 53, 55).

We sought to clarify how Nsp14 modulates diverse innate immune signaling pathways and determine whether a common mechanism may underlie the diverse effects of Nsp14. These studies demonstrate that Nsp14 not only stimulates NF-κB and ERK but also the other main arms of MAP kinase signaling, as evidenced by p38 MAP kinase, ERK and JNK activation. In addition, we demonstrate that Nsp14 downregulates not only the IFNAR1 subunit of the IFNα/β receptor but also the IFNGR1 subunit of the IFNγ receptor. In both cases, receptor downregulation occurs via a lysosomal pathway. Notably, by assessing a panel of Nsp14 point mutants and truncation mutants, we demonstrate that inactivation of N7-MTase disrupts NF-κB and p38 activation and IFNAR1/IFNGR1 downregulation and that each activity follows the same pattern of sensitivity to the mutants tested. We further demonstrate that Nsp14 interacts with the host protein Tollip, a multifunctional protein that acts as an inhibitor of NF-κB activation via Toll-like receptor and IL-1R signaling and a selective autophagy receptor that directs ubiquitinated proteins towards lysosomal degradation (56). We find that Tollip can antagonize Nsp14 augmentation of NF-κB activation upon TLR4 stimulation. Further, knockdown of Tollip counteracts the downregulation of IFNAR1 and IFNGR1 in the context of SARS-CoV-2 infection.

## Results

### SARS-CoV-2 Nsp14 activates the NF-κB and MAPK signaling pathways and induces cytokine production

We generated constructs expressing the full length Nsp14, as well as select point mutants in the exonuclease (ExoN) domain (D90A/E92A, E191A, H268A, D273A) and in the N7-methyltransferase (N7-MTase) domain (N306A, D331A) (Fig. 1A). These point mutations target enzymatic activity (14, 57, 58). We also deleted amino acids 1-60 which constitutes the Nsp10 binding site of Nsp14 (ΔNsp10bs-Nsp14) and generated separate deletions of the entire ExoN and N7-MTase domains (Fig. 1A) (58). To determine the effects of these various Nsp14 expression constructs on LPS-induced activation of NF-κB responses, reporter gene assays were performed in HEK293T cells stably expressing human TLR4, MD2 and CD14 (hTLR4-MD2-CD14). Full length, wildtype Nsp14, significantly enhanced LPS-induction of the NF-κB reporter, relative to an empty vector, untreated control (Fig. 1B). The ΔNsp10bs-Nsp14 mutant yielded comparable results, indicating that the Nsp10 binding site is dispensable for this activity. Enhancement of NF-κB induction was also either similar to wildtype or only modestly reduced for point mutations in the ExoN domain. However, constructs containing the N306A or D331A N7-MTase mutations had impaired activity, with fold increases similar to the empty vector control. Impaired enhancement of NF-κB induction was also observed for constructs expressing only the ExoN domain or only the N7-MTase domain, indicating that co-expression of both domains is required for this activity (Fig. 1B). Expression of the Nsp14 constructs was confirmed by immunoblot, with some variation in levels.

**Figure 1.**
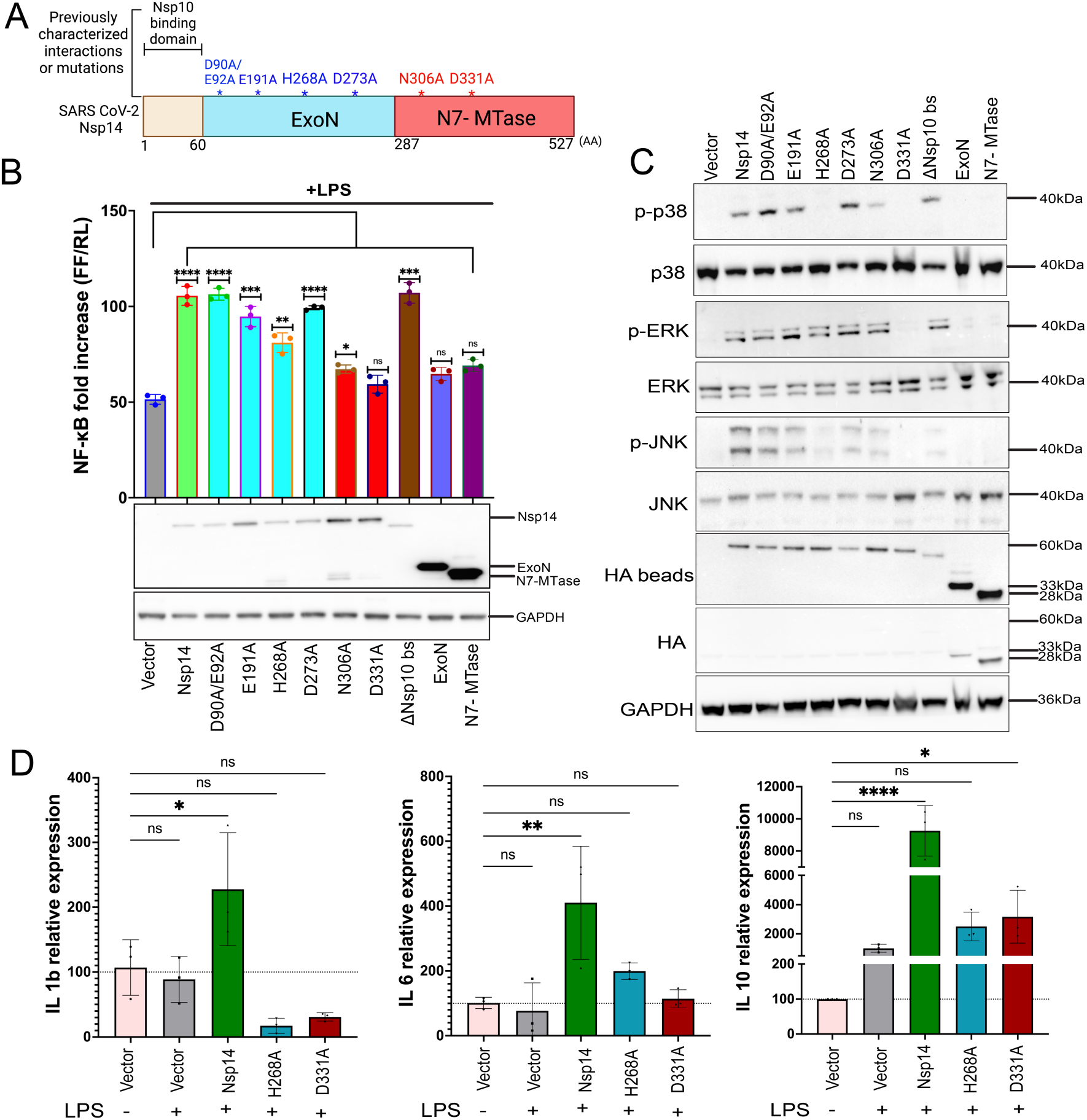
SARS CoV-2 Nsp14 activates NF-κB and MAPK signaling. A. Schematic depiction of Nsp14 protein domains, including the amino-terminal 60 amino acid residues needed for binding to Nsp10 (tan), the exonuclease (ExoN) domain (blue) and N7-methyltransferase (N7-MTase) domain (red). Amino acid residues at the boundaries of the domains are indicated at the bottom. Key amino acid residues that affect ExoN or N7-MTase activities are indicated at the top. B. HEK293 cells that express TLR4, MD2 and CD14 were transfected with an NF-κB-firefly luciferase (FF) reporter plasmid, a constitutively expressing *Renilla* luciferase (RL) and plasmids that express HA-tagged versions of the indicated proteins. At 24 h post-transfection, cells were treated with LPS. Eighteen h later a dual luciferase reporter assay was performed. The data are presented as fold induction relative to an empty vector, mock-treated control. Error bars represent mean ± SD (n = 3). One way ANOVA was used to determine statistical significance (*P* ≤ 0.05 = ∗, *P* < 0.01 = ∗∗, *P* < 0.001 = ∗∗∗, *P* < 0.0001 = ∗∗∗∗, not significant = ns). Cell lysates were analyzed by western blot. Vector, empty expression plasmid; Nsp14, full-length Nsp14; full-length Nsp14s with point mutations are indicated; ΔNsp10 bs-Nsp14, Nsp14 lacking residues 1-60; ExoN, Nsp14 residues 1-287; N7-MTase, only Nsp14 287-527. C. Immunoblots for total and phosphorylated (p) MAPK pathway proteins in the lysates of HEK293T cells transfected with empty vector or the indicated HA-tagged wildtype and mutant Nsp14s. HA beads, HA-Nsp14 proteins were concentrated by anti-HA immunoprecipitation and then analyzed by immunoblotting. D. Expression levels of IL1b, IL6 and IL10 as determined by RT-qPCR in cells transfected with empty vector (Vector), Nsp14 and the indicated Nsp14 mutants. Cells were mock-treated or treated with LPS, as indicated. Error bars represent mean ± SD (n = 3). One way ANOVA was used to determine statistical significance (*P* ≤ 0.05 = ∗, *P* < 0.01 = ∗∗, *P* < 0.0001 = ∗∗∗∗, not significant = ns).

Because SARS-CoV-2 also activates pro-inflammatory MAPK signaling (18, 19), we investigated the capacity of Nsp14 to activate the MAPK pathways. HEK293T cells were transfected with the wildtype and mutant Nsp14s and the status of MAPK family proteins in lysates was assessed by immunoblot analysis. Because HA-Nsp14 expression was difficult to detect, anti-HA immunoprecipitations were performed, and the precipitated material was assessed by immunoblotting. While total levels of p38, extracellular-signal-regulated kinase (ERK) and Jun amino-terminal kinase (JNK) were largely unaffected by Nsp14, Nsp14 expression led to an increase in phosphorylation of the three major families of MAPKs as evidence by p38, ERK and JNK phosphorylation (Fig. 1C). For ERK and JNK activation was also observed for the ΔNsp10bs mutant and for the D90A/E92A, E191A, H268A (weak for JNK) and D273A point mutants in the ExoN domain. With regard to N7-MTase mutants, some degree of activation was seen for the N306A mutant, while the D331A mutant did not stimulate the phosphorylation of ERK or JNK. One possible explanation for the different behavior of the two mutants is that N306A retained ∼30% residual activity in an in vitro enzymatic assay of N7-MTase activity, while this activity was almost completely abolished for D331A (14). The findings were very similar for p38, with induction of phosphorylation detected for most mutants but not for the N7-MTase D331A mutant. For p38, the ExoN H268A mutant also failed to induce phosphorylation, although the remaining three point mutants in ExoN retained the ability to promote phosphorylation. The individually expressed ExoN or N7-MTase domains failed to activate p38, ERK or JNK. Overall, the results indicate that Nsp14 activates both the NF-κB and MAPK signaling pathways, that neither of its major enzymatic domains when individually expressed is sufficient for the activation, and that the N7-MTase domain appears to play a critical role based on loss of activation by the catalytically dead D331A mutant.

Because activation of both the NF-κB and MAPK signaling pathways can contribute to inflammatory responses, we determined the impact of Nsp14 expression on cytokine induction in HEK293-hTLR4-MD2-CD14 cells in the presence of LPS, with transcript levels quantified by RT-qPCR. Wild-type Nsp14 enhanced expression of the typically proinflammatory cytokines IL-1β and IL-6 and the anti-inflammatory cytokine IL-10 (Fig. 1D). The induction of IL-1β, IL-6 and IL-10 by Nsp14 mutants correlated well with the NF-κB and MAPK pathway assays (Fig. 1B-C).

### SARS-CoV-2 Nsp14 downregulates expression of IFNAR1 and IFNGR1 impairing interferon signaling pathways

To assess the role of Nsp14 in IFN responses, we examined the expression of endogenous interferon receptors in cells transfected with an Nsp14-GFP fusion construct. Cells expressing Nsp14-GFP exhibited very low or negligible levels of both IFNAR1 and IFNGR1 (Fig. 2A). Levels of IFN alpha receptor 2 (IFNAR2), in contrast, appeared relatively unaffected by Nsp14-GFP (Fig. 2A). When quantified, IFNAR1 and IFNGR1 staining intensity was significantly lower for cells expressing Nsp14-GFP than for cells not expressing Nsp14-GFP, while IFNAR2 staining intensity did not differ between Nsp14-GFP-positive and –negative cells (Fig. 2B). Cultures not transfected with Nsp14-GFP displayed relatively uniform expression of all three receptors (Fig. S1).

**Figure 2.**
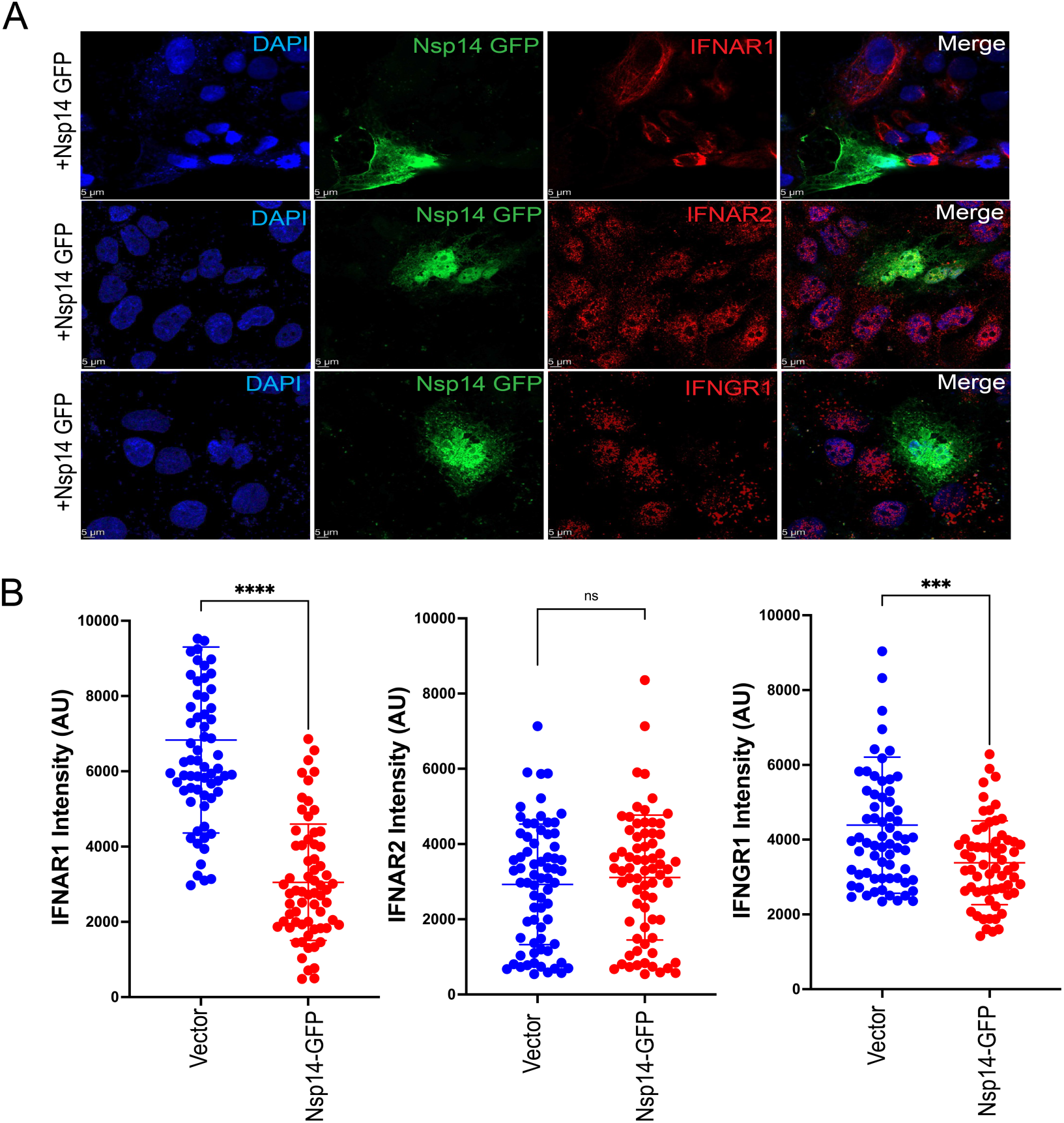
SARS CoV-2 Nsp14 downregulates the expression of IFNAR1 and IFNGR1. A. Confocal laser scanning microscopy image of interferon receptor expression levels in Huh7 cells transfected with an Nsp14-GFP expression plasmid. Blue, DAPI staining of nuclei. Green, Nsp14-GFP. Red, the indicated endogenous receptor. B. Quantification of IFNAR1, IFNAR2, IFNGR1 staining intensity in cells transfected with empty vector (Vector) or Nsp14-GFP. Each dot indicates the value for a single cell. Error bars represent mean ± SD (n = 3). Unpaired two-tailed student’s t test was used to determine statistical significance (*P* < 0.001 = ∗∗∗, *P* < 0.0001 = ∗∗∗∗, not significant = ns).

We next assessed the impact of our panel of Nsp14 constructs on IFNAR1, IFNAR2 and IFNGR1 by immunoblot. In the presence of wild-type Nsp14, endogenous levels of both IFNAR1 and IFNGR1 were substantially reduced, while levels of IFNAR2 were unaffected (Fig. 3A). Results were similar for the Nsp14 mutants tested, with the exception of the D331A mutant and the individually expressed ExoN and N7-MTase domains, which failed to reduce levels of IFNAR1 or IFNGR1. The H268A mutant exhibited an intermediate phenotype, with partial rescue of IFNAR1 levels, although levels were lower than vector control. The results mirror our findings regarding the NF-κB and MAPK signaling pathways, where both domains of the bifunctional Nsp14 protein were required to observe the phenotypes, and where the D331A point mutant, which abolishes N7-MTase activity, resulted in a complete loss of activity.

**Figure 3.**
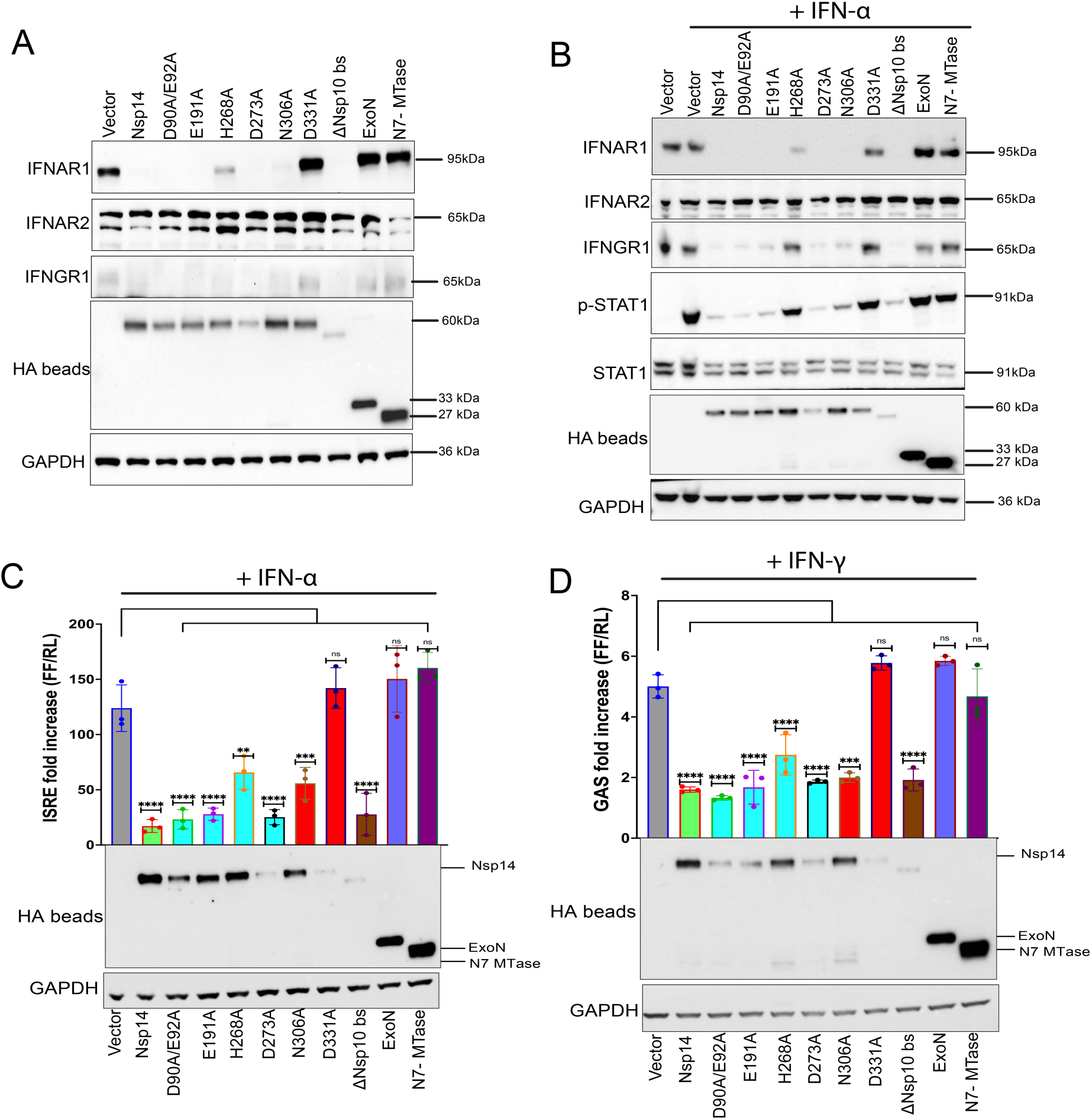
Nsp14 inhibits IFNα and IFNγ signaling via downregulation of IFNAR1 and IFNGR1. A. Immunoblots assessing levels of endogenous IFNAR1, IFNAR2, IFNGR1 and GAPDH in HEK293T cells transfected with empty vector (Vector) or plasmids expressing the indicated HA-tagged Nsp14s. HA beads, HA-Nsp14 proteins were concentrated by immunoprecipitation with beads conjugated to anti-HA antibodies and then analyzed. B. Immunoblots to assess the effect of Nsp14 on STAT1 phosphorylation 30 minutes after IFNα addition. Transfections were performed as in panel A. Blotting included antibodies to detect total and tyrosine-phosphorylated STAT1. C. Effects of transfecting the indicated expression plasmids on transcriptional responses to IFNα as assessed by reporter gene assay. HEK293T cells were transfected with an ISRE-firefly luciferase reporter plasmid, a constitutively expressing *Renilla* luciferase and the indicated plasmids. At 24 h post-transfection, cells were treated with IFNα. Twenty four hours later, luciferase activities were determined. Firefly luciferase values were normalized to *Renilla* luciferase values and data are reported as fold increase relative to empty vector, mock treated samples. Error bars represent mean ± SD (n = 3). One-way ANOVA was used to determine statistical significance (*P* < 0.01 = ∗∗, *P* < 0.001 = ∗∗∗, *P* < 0.0001 = ∗∗∗∗, not significant = ns). Cell lysates were analyzed by immunoblot. D. Effects of transfecting the indicated expression plasmids on transcriptional responses to IFNγ as assessed by reporter gene assay. Transfections were performed as in panel C except a GAS-firefly luciferase reporter plasmid was used in place of the ISRE firefly luciferase plasmid and cells were treated with 500U/ml IFN-γ. Analysis was as in C. Cell lysates were analyzed by immunoblot. Labeling of immunoblots for C and D was the same as for A.

To investigate the effects of IFN receptor dysregulation on downstream signaling, we assessed the phosphorylation status of STAT1, a key transcription factor that mediates IFNαβ signaling. We compared levels of basal and activated (tyrosine phosphorylated) STAT1 in lysates of cells transfected with vector alone without IFN treatment; as well as cells transfected with vector alone, with the various Nsp14 constructs in the presence of universal type I IFN. Thirty minutes post IFN addition, basal STAT1 levels were similar for all proteins tested (Fig. 3B). As expected, IFN treatment resulted in the appearance of tyrosine phosphorylated STAT1 (p-STAT1) in the vector control. Consistent with the observed downregulation of IFNAR1, levels of p-STAT1 were diminished in the presence of wild-type Nsp14 (Fig. 3B). Similar reductions in p-STAT1 were observed for the Nsp14 mutants, with the exceptions of H268A, D331A, and the individual ExoN and N7-MTase domains, all of which displayed activation similar to controls. In addition, as was the case in the absence of IFN (Fig. 3A), both IFNAR1 and IFNGR1 levels were substantially reduced in the presence of IFN treatment for cells expressing wild-type Nsp14 (Fig. 3B). This was also true for the Nsp14 mutants, again with the exception of H268A, D331A, and the individual ExoN and N7-MTase domains, all of which were substantially impaired in the ability to downregulate IFNAR1 and IFNGR1. Finally, as seen in the absence of IFN treatment, IFNAR2 levels were unaffected by Nsp14 in the presence of IFN, indicating a specificity to dysregulation of receptor levels (Fig. 3B).

To further define the impact of receptor downregulation on type I- and type II-IFN-mediated signaling, interferon-stimulated response element (ISRE)-firefly luciferase reporter gene assays were performed in the absence or presence of wild-type or mutant Nsp14s. As expected, given the reduced IFNAR1 levels and loss of STAT1 activation, Nsp14 repressed the activation of the ISRE reporter in the presence of type I IFN (Fig. 3C). This repression was relieved for the D331A mutant and for the ExoN and N7-MTase individual domain mutants (Fig. 3C). Similar findings were obtained using cells transfected with an interferon gamma-activated sequence (GAS)-firefly luciferase reporter plasmid and treated with IFNγ (Fig. 3D). Expression of each Nsp14 construct was confirmed by immunoblot analysis for these studies (Fig. 3C-D).

### Nsp14 downregulation of IFNAR1 and IFNGR1 involves a lysosomal pathway

Expression of Nsp14 has been shown to induce widespread transcriptional changes resembling those following SARS-CoV-2 infection, effects that require the N7-MTase domain of the protein (45). Further, SARS-CoV-2 Nsp14 has been identified as an inhibitor of nuclear mRNA processing and export (44). To investigate whether Nsp14 modulates interferon receptor expression at the level of transcription, the mRNA levels of IFNAR1, IFNAR2, and IFNGR1 were quantified by RT-qPCR in cells transfected with empty vector or the different Nsp14 constructs. Transcript levels of IFNAR1 and IFNGR1 were similar for cells expressing wild-type Nsp14, H268A or D331A, as compared with the vector control (Fig. 4A). Despite not exhibiting obvious differences at the protein level, IFNAR2 transcript levels were modestly reduced to varying degrees with expression of wild-type Nsp14 or the D331A or H268A mutants (Fig. 4A). These results indicate that Nsp14 downregulation of IFNAR1 and IFNGR1 expression occurs at a post-transcriptional level. A prior study found that Nsp14 inhibition of translation was abrogated by mutations that disrupt ExoN or N7-MTase activities (43). Our ExoN mutant data therefore suggests that translation inhibition does not explain IFNAR1 and IFNGR1 downregulation.

**Figure 4.**
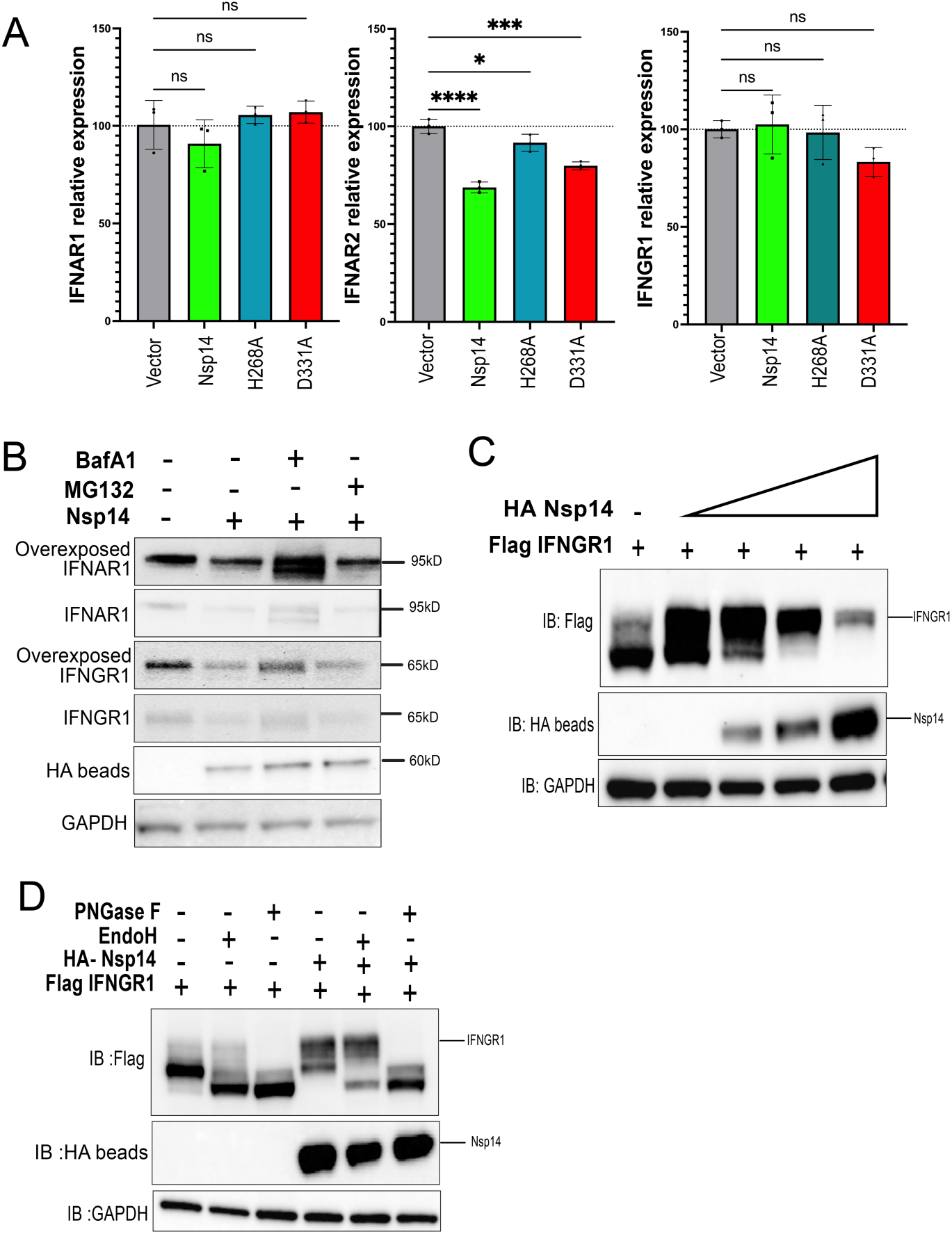
Mechanisms of IFNAR1 and IFNGR1 downregulation. A. RT-qPCR analysis of IFNAR1, IFNAR2, IFNGR1 mRNA expression levels in Huh7 cells 24 h post-transfection with empty vector (Vector), Nsp14 and the indicated Nsp14 mutants. Error bars represent mean ± SD (n = 3). One way ANOVA was used to determine statistical significance (*P* ≤ 0.05 = ∗, *P* < 0.001 = ∗∗∗, *P* < 0.0001 = ∗∗∗∗, not significant = ns). B. Immunoblot for endogenous level of IFNAR1 and IFNGR1 in HEK293T cells overexpressing HA-Nsp14 in the absence or presence of BafA1 (2 µM) or MG132 (50 µM) for 6 hr before harvesting. HA bead, samples were concentrated by immunoprecipitation with anti-HA antibody conjugated beads prior to immunoblotting. GAPDH served as a loading control. C. Immunoblot demonstrating effects of increasing concentrations of Nsp14 on overexpressed IFNGR1 levels and migration on SDS-PAGE. HA beads, samples were concentrated by immunoprecipitation with anti-HA antibody conjugated beads prior to immunoblotting. D. Immunoblot of cell lysates from HEK293T overexpressing Flag-IFNGR1 and HA-Nsp14 that were mock treated or treated with PNGase F or Endo H. GAPDH served as a loading control. HA bead, samples were concentrated by immunoprecipitation with anti-HA antibody conjugated beads prior to immunoblotting.

To address the mechanism of this post-transcriptional and post-translational regulation of IFN receptor levels, we examined the impact of inhibitors of the proteasomal and lysosomal pathways. The Nsp14-dependent reduction in endogenous levels of IFNAR1 and IFNGR1 was maintained in the presence of the proteasome inhibitor MG132, but was diminished in the presence of the lysosomal inhibitor bafilomycin A1 (BafA1), which rescued both IFNAR1 and IFNGR1 levels (Fig. 4B). Expression of increasing amounts of HA-Nsp14 resulted in a dose-dependent reduction in the expression of co-transfected Flag-IFNGR1 (Fig. 4C). On Western blot analysis, at least two bands were observed for affinity-tagged, over-expressed IFNGR1, with the faster-migrating band most affected by Nsp14 expression (Fig. 4C). To determine whether the multiple bands might represent differentially glycosylated forms, we subjected the lysates to digestion with peptide:N-glycosidase F (PNGase F), which removes almost all types of *N*-linked (Asn-linked) glycosylation and with Endoglycosidase H (Endo H), which more selectively removes high mannose and certain hybrid types of *N*-linked carbohydrates. In the absence of Nsp14, Flag-IFNGR1 was susceptible to digestion by both PNGase F and Endo H, with a shift in the band to the faster-migrating form (Fig. 4D). With co-expression of Nsp14, Flag-IFNGR1 remained susceptible to digestion by PNGase F but displayed substantial resistance to Endo H (Fig. 4D). This suggests that Nsp14 selectively promotes degradation of Flag-IFNGR1 forms in the ER and in early regions of the Golgi complex, allowing more Endo H-resistant forms to predominate in its presence.

### Identification of the autophagy selective receptor protein Tollip as an interacting partner of SARS-CoV-2 Nsp14

Tollip is an adaptor protein expressed in numerous cell types that plays roles in diverse intracellular signaling pathways. Among its functions, Tollip mediates delivery of cargo proteins to the lysosome for degradation, including the receptors for IL-1 and TGFβ (59, 60). Tollip has also been implicated in the selective clearance of aberrant ER cargos (61). Ongoing work focused on Tollip identified an interaction between Nsp14 and Tollip. This is illustrated by immunofluorescence analysis in which Nsp14 colocalizes with endogenous Tollip (Fig. 5A). Coimmunoprecipitation analysis also demonstrated interaction between HA-tagged Nsp14 and endogenous Tollip. Although not precipitated with the empty vector control, Tollip co-precipitated with wild-type Nsp14 and also with the individual ExoN and N7-MTase domains (Fig. 5B).

**Figure 5.**
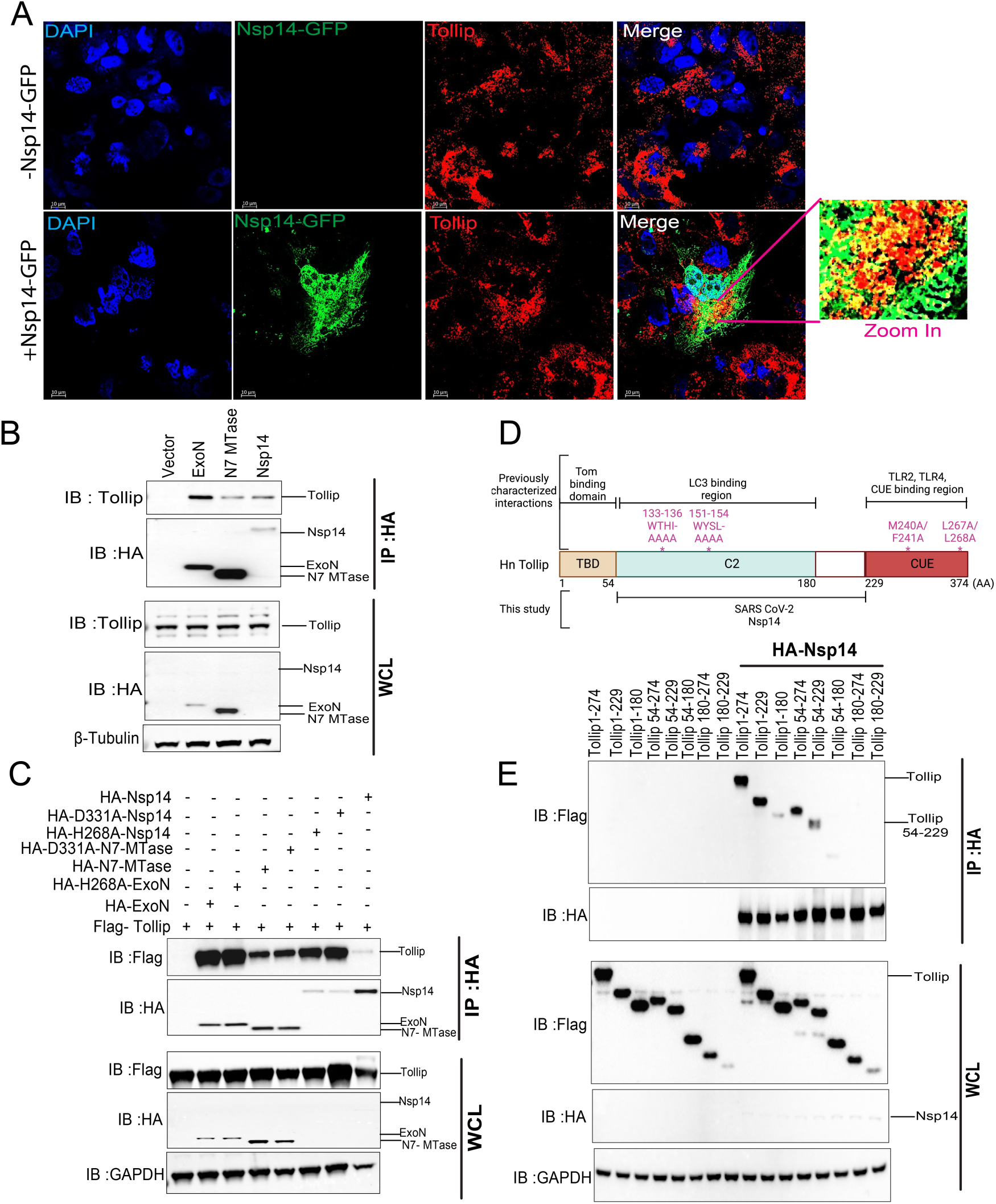
SARS-CoV-2 Nsp14 co-localizes and interacts with Tollip. A. Immunofluorescence to assess localization of Nsp14-GFP and endogenous Tollip in Huh7 cells. Blue, DAPI (nuclei); Green, Nsp14-GFP; Red, Tollip. B. Co-IP of endogenous Tollip with transfected empty vector (Vector), HA-Nsp14 ExoN domain, N7 MTase domain or full-length Nsp14. Immunoblots of the immunoprecipitations (IP:HA) and whole cell lysates (WCL) are shown. Blots were probed with anti-Tollip, anti-HA and anti-GAPDH antibodies, as indicated. C. Interaction of Flag-Tollip and the indicated HA-tagged Nsp14 constructs was assessed by coIP. IP:HA, samples subjected to anti-HA immunoprecipitation. Immunoblots of the immunoprecipitations (IP:HA) and whole cell lysates (WCL) are shown. Blots were probed with anti-Flag, anti-HA and anti-GAPDH antibodies, as indicated. D. Schematic depiction of Tollip protein domains and key residues for catalytic activity and ubiquitin binding. TBD, Tom1 binding domain; C2, central conserved 2 (C2) domain involved in Ca^2+^-dependent membrane-targeting, LC3 binding; CUE, coupling of ubiquitin to ER degradation domain. Amino acid residues at the boundaries of domains are indicated below the diagram. Point mutants used in this study are indicated above the diagram. The region that interacts with Nsp14, as determined by this study, is indicated below the diagram. E. Domain mapping studies. HEK293T cells were co-transfected with empty vector or HA-Nsp14 and the indicated Flag-tagged Tollip constructs and anti-HA immunoprecipitations were performed. Immunoblots of the immunoprecipitations (IP:HA) and whole cell lysates (WCL) are shown. Blots were probed with anti-Flag, anti-HA and anti-GAPDH antibodies, as indicated.

Because Tollip interacts with both the Nsp14 ExoN and MTase domains, we tested whether the catalytic activities of the domains were required for the interaction. In addition to the previously described H268A and D331A mutations in full-length Nsp14, we also introduced these mutations into the truncated ExoN and N7-MTase domains, respectively. The HA-tagged Nsp14 wild-type or mutants were co-expressed with Flag-tagged Tollip in HEK293T cells and immunoprecipitations were performed with anti-HA beads. Interaction with Tollip was maintained for both the exonuclease mutant (H268A) and the N7-MTase mutant (D331A), whether introduced into individual domains or into the full length Nsp14 (Fig. 5C), indicating that Nsp14 catalytic activities are dispensable for the Tollip interaction.

Tollip is a modular protein, containing an N-terminal Tom1-binding domain (TBD) involved in endosomal sorting, a central conserved 2 (C2) domain which interacts with phosphoinositides and LC3, and a C-terminal CUE domain involved in ubiquitin-binding (Fig. 5D) (56, 62, 63). To determine which of these domains are important for Nsp14 interaction, we generated a series of truncation mutants. Flag-tagged Tollip constructs were co-expressed with HA-tagged Nsp14. When HA-Nsp14 was immunoprecipitated, the Tollip domain spanning amino acids 54-229 or constructs in which this region was retained, were co-precipitated (Fig 5E). The Tollip 1-180 and 54-180 constructs co-precipitated very weakly suggesting that 54-180, which corresponds to the C2 domain, may be sufficient for interaction. However, because 54-229 clearly binds better, residues 180-229 contribute to a strong interaction. Together, the findings emphasize the importance of the C2 domain as well as sequences immediately C-terminal to the C2 domain in the interaction.

### Tollip partially reverses Nsp14 mediated NF-κB activation and cytokine expression

Tollip regulates several important inflammatory pathways and has been shown to inhibit NF-κB signaling triggered by IL-1β and by toll-like receptors TLR2 and TLR4 (64–66). Given this and the interaction between Tollip and Nsp14, we asked whether Tollip expression would affect the Nsp14-mediated activation of NF-κB. In agreement with our prior findings (Fig. 1B), expression of Nsp14 led to a significant increase in LPS-stimulated NF-κB reporter activity; this increase was found to be substantially dampened by the overexpression of Tollip (Fig. 6A). When Tollip was expressed at high levels in the presence of Nsp14, NF-κB reporter activity was reduced to that of control cells transfected with empty vector alone. Similarly, expression of Nsp14 increased the LPS-induced expression of IL-1β and IL-8 transcripts but co-expressed Tollip reversed these effects (Fig.6B).

**Figure 6.**
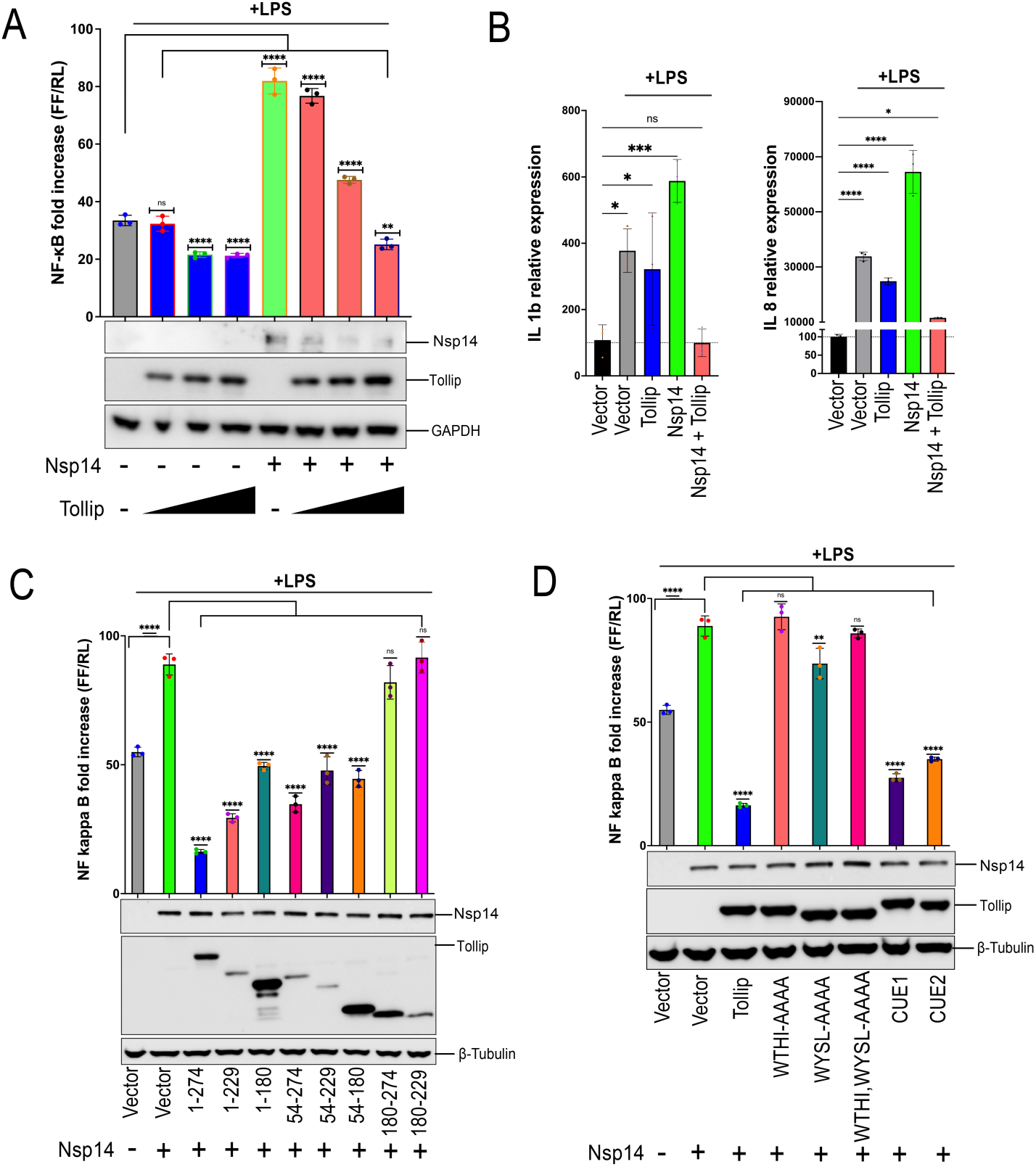
Tollip counteracts Nsp14-mediated NF-κB activation. A. Effect of Tollip levels on Nsp14 activation of NF-κB. HEK293 cells that express TLR4, MD2 and CD14 were transfected with an NF-κB-firefly luciferase (FF) reporter plasmid, a constitutively expressing *Renilla* luciferase (RL), HA-Nsp14 and increasing amounts of Flag-tagged Tollip. Eighteen h later a dual luciferase reporter assay was performed. Firefly luciferase activity was normalized to *Renilla* luciferase activity. The data are presented as fold induction relative to an empty vector, mock-treated control. Error bars represent mean ± SD (n = 3). One-way ANOVA was used to determine statistical significance (*P* < 0.01 = ∗∗, *P* < 0.0001 = ∗∗∗∗, not significant = ns). Cell lysates were analyzed by immunoblot for the indicated proteins. B. IL1β and IL8 mRNA levels in in HEK293 cells that express TLR4, MD2 and CD14 transfected with empty vector (Vector), Tollip and/or Nsp14 plasmids, in the absence or presence of LPS, as indicated. Error bars represent mean ± SD (n = 3). One way ANOVA was used to determine statistical significance (*P* ≤ 0.05 = ∗, *P* < 0.001 = ∗∗∗, *P* < 0.0001 = ∗∗∗∗, not significant = ns). C. Domain mapping studies. HEK293-hTLR4-MD2-CD14 cells were transfected with an NF-κB-firefly luciferase reporter plasmid and the indicated expression plasmids. The absence or presence of Nsp14 expression plasmids is indicated at the bottom. At 24 h post-transfection, cells were treated with LPS and a dual luciferase reporter assay was performed the following day. Firefly luciferase activity was normalized to *Renilla* luciferase activity. Error bars represent mean ± SD (n = 3). One way ANOVA was used to determine statistical significance (*P* < 0.0001 = ∗∗∗∗, not significant = ns). Cell lysates were analyzed by western blot. D. Tollip point mutant studies. An NF-κB-firefly luciferase reporter assay was performed as above. The indicated protein expression plasmids were assayed. Error bars represent mean ± SD (n = 3). One-way ANOVA was used to determine statistical significance (*P* < 0.01 = ∗∗, *P* < 0.0001 = ∗∗∗∗, not significant = ns). Cell lysates were analyzed by western blot.

We next mapped this inhibitory activity using a series of Tollip truncation mutants. As before, Nsp14 enhanced the activity of an NF-κB reporter in the presence of LPS stimulation and full-length Tollip was inhibitory (Fig. 6C). The Tollip 1-229 mutant, which contains the TBD and the C2 domain and as well as some additional C-terminal sequence but lacks the CUE domain, retained substantial inhibitory activity (Fig. 6C). This was also true for the 54-274 mutant that lacks the TBD but contains both the C2 and CUE domains (Fig. 6C). Mutants 180-274 and 180-229, both of which lack the C2 domain, displayed little, if any, activity (Fig. 6C). Mutant 1-180, which contains the C2 domain but not additional C-terminal sequence, as well as mutants 54-229 and 54-180 showed similar and intermediate inhibitory activity (Fig. 6C). Together, the results support the importance of the C2 domain and C-terminal sequences adjacent to it and indicate that multiple regions of the protein may be required for full activity, as none of the truncation mutants are as active as full-length Tollip. We also examined the Tollip deletion mutants in the presence of LPS but in the absence of Nsp14. Overall, a similar pattern of inhibitory activities was noted, with the 1-229 and 54-274 mutants again showing the greatest activity, although the 1-180 mutant was inactive in this context (Fig. S2).

We also evaluated Tollip mutants with impaired LC3 and ubiquitin binding. We generated Tollip alanine substitution mutations (1) in the two LC3-interacting motifs of the C2 domain, both separately and combined: W133A, T134A, H135A, I136A (WTHI-AAAA); W151A, Y152A, S153A, L154A (WYSL-AAAA); and W133A, T134A, H135A, I136A, W151A, Y152A, S153A, L154A (WTHI-AAAA/ WYSL-AAAA) and (2) in the two ubiquitin-binding motifs of the CUE domain: M240A/F241A (CUE1) and L267A/L268A (CUE2) (Fig. 6D)(67, 68). Using the same reporter assay as above, in the presence of Nsp14, the LC3 binding mutants were unable to inhibit LPS-induced NF-κB reporter activity, while the ubiquitin binding mutants still inhibited (Fig. 6D), indicating that association with LC3 but not with ubiquitin may be important for Tollip’s antagonism of Nsp14. In the absence of Nsp14, a similar pattern was observed, although only partial loss of activity was seen with two of the LC3 binding mutants (Fig. S2).

### IFNAR1 and IFNGR1 levels are reduced in the context of SARS-CoV-2 infection and this reduction is blunted when Tollip expression is silenced

To address the effects of SARS-CoV-2 infection on IFNAR1 and IFNGR1 and the potential role for Tollip, we examined endogenous levels of interferon receptors in A549 cells transduced with angiotensin converting enzyme 2 (A549-ACE2), in the absence and presence of infection with SARS-CoV-2 at a multiplicity of one. In the absence of infection, IFNAR1 levels were stable between 12-36 hours post infection. After infection with SARS-CoV-2, viral N protein levels increased as expected. A reduction in IFNAR1 levels was apparent at 24 hours and levels decreased further at 36 hours post-infection (Fig. 7A). Levels of IFNGR1 were also diminished at 24 and 36 hours after infection when compared with the 0 hour time point (Fig. 7A). In this case, for unclear reasons, the initial time point showed a higher level of IFNGR1 expression in the context of infection as compared with uninfected control cells. In contrast to IFNAR1 and IFNGR1, levels of IFNAR2 remained essentially constant throughout the time course of infection and were similar to levels in uninfected controls at each time point. Further, the GAPDH levels were similar at all time points. This decrease of interferon receptor levels in the context of SARS-CoV-2 infection mirrors our findings following transfection of Nsp14 (Fig. 3 A-B). Inhibition of lysosomal degradation by BafA1 partially rescued levels of IFNAR1 and fully rescued levels of IFNGR1 (Fig. 7B), pointing to a role for lysosomal activity in the receptor downregulation, while treatment with proteasome inhibitor MG132 had no effect. This also agrees with our findings in cells transfected with Nsp14 (Fig. 4B).

**Figure 7.**
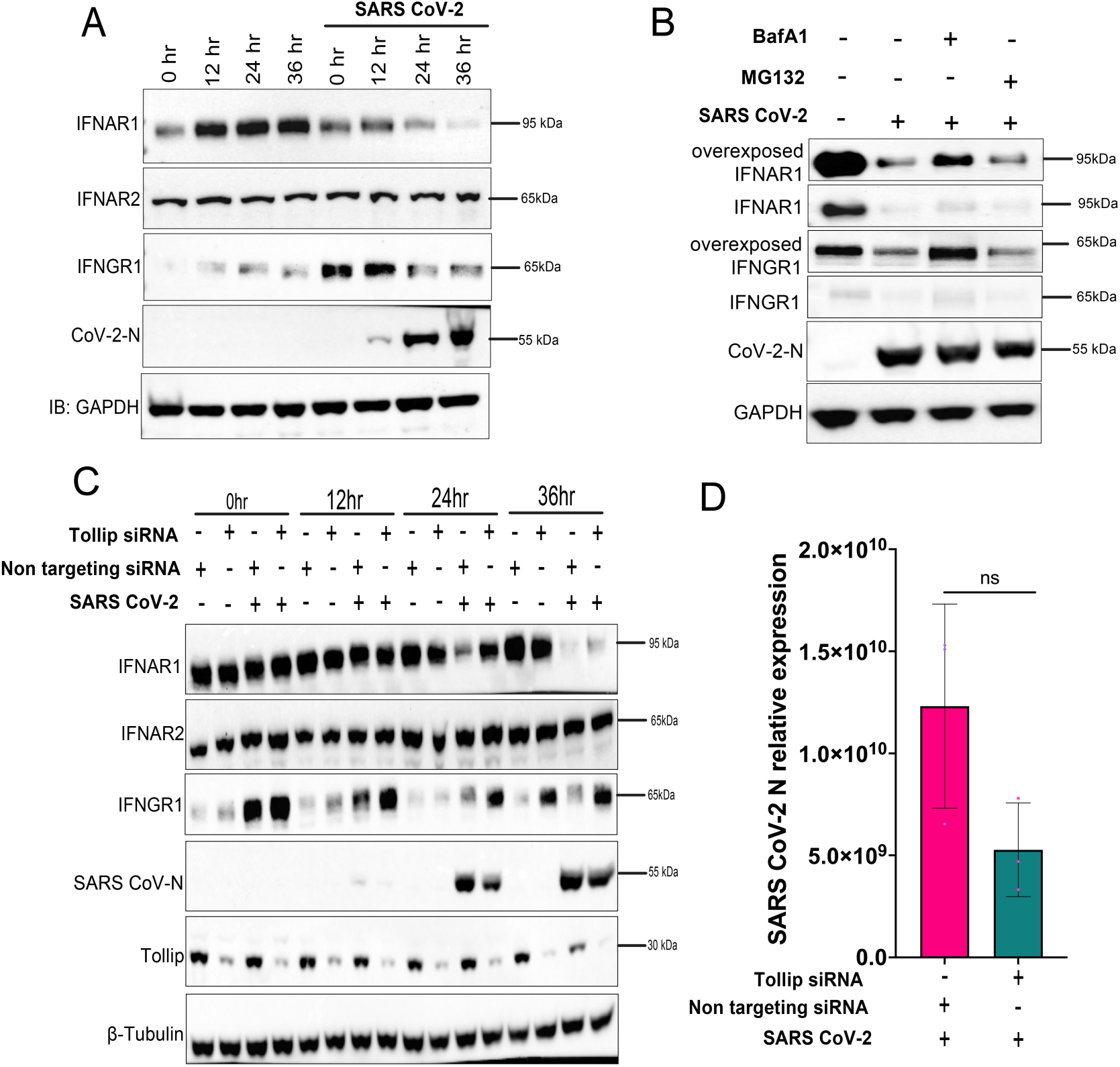
Selective downregulation of interferon receptor expression during SARS-CoV-2 infection with and without Tollip knockdown. A. Effects of SARS-CoV-2 infection on endogenous IFNAR1, IFNAR2 and IFNGR1 levels. A549-ACE2 cells were mock-infected or infected with SARS-CoV-2 (MOI=1). Lysates for immunoblotting were prepared at the indicated time points. Blots were probed with antibodies for IFNAR1, IFNAR2, IFNGR1, SARS-CoV-2 N and GAPDH, as indicated. B. Immunoblot for endogenous IFNAR1 and IFNGR1 following mock-infection or SARS-CoV-2 infection for 18h at MOI=1. As indicated, infected cells were treated with BafA1 (2 µM) or MG132 (50 µM) for 6 hr before harvesting. GAPDH was included as a loading control. C. Immunoblot analysis of A549-ACE2 cells transfected with non-targeting or Tollip specific siRNA and infected with SARS-CoV-2 (MOI=1) for indicated times post infection. Blots were probed with antibodies for IFNAR1, IFNAR2, IFNGR1, SARS-CoV-2 N and GAPDH, as indicated. D. RT-qPCR analysis of SARS-CoV-2 N mRNA expression level in A549-ACE2 transfected with non-targeting or Tollip specific siRNA and infected with SARS-CoV-2 (MOI =1). N mRNA levels were normalized to β-actin mRNA levels.

In order to evaluate the role of Tollip in interferon receptor downregulation mediated by SARS-CoV-2, we employed siRNA-mediated knockdown of Tollip expression in A549-ACE2 cells, followed by infection with SARS CoV-2. Control cells were treated with a non-targeting negative control siRNA. Depletion of Tollip impaired IFNAR1 and IFNGR1 downregulation in the setting of infection, most evidently at 24 and 36 hours post-infection (Fig. 7C). At the later time point, when Tollip levels were reduced, levels of IFNAR1 and IFNGR1 increased relative to the non-targeting control knockdown samples but remained lower than at the time of infection. IFNAR2 levels were not significantly impacted by Tollip depletion. We noticed a modest reduction in SARS-CoV-2 N protein levels on Western blot at 24 h post-infection in the setting of Tollip knockdown (Fig. 7C). Therefore, we performed RT-qPCR to examine transcript levels of the N gene at 24 hrs post-infection and observed only a modest decrease in the presence of Tollip knockdown as compared with the control and the difference did not reach statistical significance (Fig. 7D). Overall, these data suggest a role for Tollip in downregulation of IFNAR1 and IFNGR1 in the context of SARS-CoV-2 infection.

## Discussion

Our data demonstrate that full-length Nsp14 and a mutant lacking the N-terminal Nsp10 binding domain are able to stimulate NF-κB responses to LPS. Likewise, point mutants previously reported to disrupt Nsp14 ExoN activity retained the capacity to augment NF-kB activation. However, mutations that impair N7-MTase activity lost the enhancing activity, suggesting that this enzymatic function may be needed for NF-κB stimulation. Expression of either of the individual ExoN or MTase domains was also insufficient to stimulate NF-kB responses. The latter observations could be an additional reflection of the need for N7-MTase activity. That is, for SARS-CoV Nsp14, the ExoN domain, although not ExoN enzymatic activity, was found to be required for N7-MTase activity (69). Consistent with our data, other studies demonstrated that NF-κB activation was abrogated by mutations that disrupt Nsp14 methyltransferase activity but not mutants that prevent exonuclease activity (45, 47, 48). Further, expression of the ExoN or MTase domains was not sufficient to activate NF-κB (46, 48, 50).

Interestingly, our MAPK data generally paralleled the NF-κB data with full-length wildtype Nsp14, or Nsp14 deleted for the N-terminal Nsp10 binding domain activating all three of the main MAPK families (70), as demonstrated by p38 MAPK, ERK and JNK phosphorylation. Most ExoN mutants triggered p38, ERK and JNK phosphorylation. However, the H268A mutant led to ERK phosphorylation and modest but detectable JNK phosphorylation, but p38 MAPK phosphorylation was not detected. Of note, this mutant is also modestly less potent for NF-κB activation. Regardless, the data with the other ExoN mutants suggest that Nsp14 exonuclease activity is not required for MAPK activation. N7-MTase mutant D331A did not detectably activate MAPK signaling, while N7-MTase mutant N306A, which retains some N7-MTase activity in the context of SARS-CoV Nsp14 (14), was significantly impaired in this regard. This suggests that the MTase domain and perhaps MTase activity contributes to MAPK activation. As with the NF-κB assays, expression of the individual ExoN or MTase domains did not trigger MAPK signaling.

While our manuscript was in preparation, a publication reported Nsp14 activation specifically of ERK and triggering of AP-1-dependent gene expression (52). In contrast with our findings, this study reported a lack of Nsp14 activation of p38 and JNK phosphorylation. This could possibly be explained by the relatively high basal levels of phospho-p38 and phospho-JNK in the assays of Li et al. (52). Another notable difference was the capacity of the ExoN domain, expressed alone, to trigger ERK phosphorylation (52). Why this was seen in one study but not ours is unclear. However, our extensive assessment of multiple ExoN point mutants strongly suggests that Nsp14 exonuclease activity is not critical for the activation of MAPK signaling. Activation of ERK was attributed to Nsp14 interaction with the ERK kinase MEK (52). However, the broad activation of both NF-κB and MAPK signaling by wildtype Nsp14 and the parallel activities of our panel of Nsp14 mutants suggests a less pathway specific mechanism of action.

IFN responses to SARS-CoV-2, mechanisms of viral IFN evasion and the potential dysregulation of these responses in the setting of severe COVID-19 disease have been a subject of intense interest (23, 71–74). Nsp14 has been implicated in inhibition of several aspects of the IFN response, including RIG-I mediated induction of the IFNβ promoter and IRF3 nuclear accumulation (53, 54). Nsp14 has also been shown to antagonize type I IFN signaling and on immunoblot analysis to downregulate expression of the IFNAR1 chain of the IFNα receptor, possibly through lysosomal degradation (36, 53, 55). Our data confirm that Nsp14 can block type I IFN signaling through downregulation of IFNAR1. We also demonstrated no effect on IFNAR2. This downregulation is sufficient to block IFNα-induced STAT1 phosphorylation and ISG upregulation. Another finding was the downregulation of IFNGR1 by Nsp14. Treatment of cells with BafA1, an inhibitor of autophagosome-lysosome fusion and lysosome acidification (75), partially rescues both IFNAR1 and IFNGR1 from downregulation by Nsp14. This is consistent with an earlier study that suggested Nsp14 decreases levels of IFNAR1 by promoting degradation via the lysosome (55) and extends the observation to IFNGR1. Interestingly, our studies with Endo H and PNGase F digestion, which were performed in the presence of limiting amounts of Nsp14 as well as transfected IFNGR1 to prevent complete degradation of IFNGR1, suggest that Endo H sensitive, less mature forms of IFNGR1 are preferentially targeted by Nsp14, at least when IFNGR1 is over-expressed. These are likely forms present in the ER or early regions of the Golgi complex.

We did not test the impact of SARS-CoV-2 infection on NF-κB and MAPK signaling, because activation of these pathways has been previously demonstrated (17–19, 76). We did assess the effects of SARS-CoV-2 infection on IFN receptors. Importantly, the effects of infection paralleled those of Nsp14 alone. Over the course of infection, IFNAR1 and IFNGR1 decreased, whereas IFNAR2 remained stable. Of note, after incubation of virus with the cells, the levels of IFNGR1 detected by immunoblot were higher than for the uninfected cells. The basis for this increase is not clear, but the decrease in IFNGR1 over the course of infection was evident. In addition, as in the transfection studies, BafA1 but not MG132 partly restored IFNAR1 and IFNGR1 levels. These parallels support the relevance of Nsp14 receptor downregulation for SARS-CoV-2 infection. It remains to be determined how broadly/selectively Nsp14 affects cell surface receptors.

Tollip, which we serendipitously found interacts with Nsp14 in co-IP assays, modulates TLR4 signaling and promotes lysosomal degradation of a variety of proteins. Therefore, it was of interest to explore the basis of Nsp14-Tollip interaction and to what extent Tollip can influence the immune modulatory properties of Nsp14. Prior studies demonstrated that Tollip over-expression inhibits IL-1R, TLR4 or TLR2 mediated NF-κB activation (60, 64, 65, 77). Described mechanisms of Tollip inhibition include binding to the TIR domains of these receptors and binding to the kinase IRAK1 (64, 65). Studies that sought to define the TLR4 interacting regions of Tollip implicated the C2 domain as well as more C-terminal sequences (65). This roughly mirrors the regions that interact with Nsp14. Our data with Tollip point mutants implicate phosphoinositide/LC3 binding in inhibition of Nsp14-mediated NF-κB activation and in NF-κB inhibition in the absence of Nsp14. Our data with the CUE1 and CUE2 mutants indicate that ubiquitin binding is not needed for Tollip inhibition of Nsp14-mediated or Nsp14-independent NF-κB activation. Prior studies have defined mutations in the C2 domain, including mutation K150E that disrupt binding to phosphoinositides and LC3 (67, 78). Notably, K150 is adjacent to the to the LC3 binding mutant WYSL-AAAA that we tested, and K150E was previously demonstrated to exhibited reduced capacity to block LPS signaling (78). Overall, the similarities between inhibition in the absence and presence of Nsp14 suggest that Tollip-Nsp14 binding may not fully account for the effects of Tollip on Nsp14-NF-κB stimulation.

Tollip also regulates trafficking of a variety of membrane-associated receptors, suggesting possible relevance to Nsp14 downregulation of IFNAR1 and IFNGR1. For example, IL-1β triggers the ubiquitination of Interleukin-1 Receptor Type 1 (IL-1R1) and this allows interaction with Tollip via its ubiquitin binding CUE domain, leading to the trafficking of IL-1R1 to late endosomes for degradation (60). Tollip also promotes the degradation of TGF-β type I receptor (TβRI) (59) and directs select aberrant membrane proteins in the ER for degradation via the lysosome (61). Suggesting that Tollip is relevant to the Nsp14-mediated downregulation of IFNAR1 and IFNGR1, siRNA knockdown of Tollip in the context of SARS-CoV-2 infection reduced the loss of IFNAR1 and IFNGR1. Rescue of IFNGR1 levels was more apparent but, for IFNAR1, Tollip knockdown had a clear effect at 24 hours post-infection and a more modest effect was evident at 36 hours post infection. A prior study reported that Tollip promotes ACE2 degradation and impairs SARS-CoV-2 infection (79). Therefore, it was important to assess the effects of Tollip knockdown in our assays. We did not detect enhanced infection upon Tollip knockdown. This may reflect saturating levels of ACE2 in the A549 cells engineered to express ACE2 that we used for these studies. Interestingly, we detected a modest decrease in viral N protein levels that was most evident at 24 hours post-infection. This corresponded to a modest but not statistically significant decrease in viral RNA. While we cannot exclude some impact of this impaired viral gene expression on IFNAR1 and IFNGR1 downregulation, the effects are sufficiently modest that they likely do not explain the effects of Tollip knockdown.

Cumulatively, our data highlight the contrasting effects of Nsp14, which result in pro-inflammatory innate immune pathway activation and simultaneous downregulation of IFNα and IFNγ signaling. It is striking that the anti-IFN receptor activities of our Nsp14 mutants paralleled the effects on NF-κB and MAPK signaling, such that activators of NF-kB and MAPK are down-regulators of IFNAR1 and IFNGR1. Although the underlying mechanism remains unclear, this suggests a common activity of Nsp14 underpinning these diverse effects. Prior studies have demonstrated NF-kB activation by Nsp14 (46, 48, 50). NF-κB activation has been variously attributed to interaction of Nsp14 with IMPDH2 (45, 46) and with the IKKα, IKKβ, NEMO complex (47–49). Nsp14 may undergo linear ubiquitination to recruit IKKα, IKKβ and NEMO (50). How these observations relate to effects on MAP kinase activation of IFNAR1/IFNGR1 downregulation remains to be determined.

With regard to Tollip, while it can counteract the effects of Nsp14 on NF-κB activation, analysis of Tollip mutants does not clearly distinguish between Tollip-Nsp14 interaction versus more general effects of Tollip on NF-κB signaling as a mechanism of inhibition. In the context of SARS-CoV-2 infection, the presence of Tollip contributes to IFNAR1 and IFNGR1 downregulation, suggesting a possible role of the Nsp14-Tollip interaction in receptor downregulation. As with NF-κB activation, it remains to be determined whether and how Nsp14-host protein interactions relate to the effects of N7-MTase mutations. Finally, it seems probable that the mechanisms that activate NF-κB and MAPK and promote IFNAR1 and IFNGR1 downregulation should impact other pathways. The full breadth of effects of Nsp14 on cellular signaling pathways warrants further analysis.

## Materials and Methods

### Cell lines and viruses

HEK293T (ATCC, CRL-3216), HEK293/hTLR4A-MD2-CD14 (Invivogen, 293-htlr4md2cd14), Huh7 (a generous gift from the Gordan laboratory at the University of California at San Francisco), A549-ACE2 (a generous gift from the García-Sastre laboratory at the Icahn School of Medicine at Mount Sinai), and VeroE6 (ATCC, CCL-81 ™) cells were maintained in Dulbecco’s modified Eagle’s medium (Corning) with 10% fetal bovine serum (Gibco) at 37°C in a humidified atmosphere with 5% CO_2_. Where indicated in the Results, cells were treated with Bafilomycin A1 (Invivogen tlrl-baf1), MG132 (Sigma-Aldrich M7449), LPS (Invivogen tlrl-pb5lps), human IFN-γ (PeproTech 300-02-100ug) and Universal^TM^ Type I IFN (Human IFN-Alpha Hybrid Protein) (PBL Assay Science 11200-1).

SARS-CoV-2 isolate USA-WA1/2020 was kindly provided by the García-Sastre laboratory at Icahn School of Medicine at Mount Sinai. Virus stocks were prepared on Vero E6 cells. Viruses were titered by plaque assay and stored at −80 °C. The human cell line A549 transduced with angiotensin-converting enzyme 2 (A549-ACE2) was infected at the indicated MOIs with SARS-CoV-2.

### Plasmids

Cloned SARS-CoV-2 Nsp14 (a kind gift from the Krogan laboratory at the University of California at San Francisco) was used as a template to amplify by PCR Nsp14 wildtype and mutants D90A/E92A, H268A, D273A, N306A, D331A, ΔNsp10bs, ExoN, N7-MTase. These were cloned with an N-terminal HA tag into the mammalian expression plasmid pCAGGS. Nsp14 with GFP tag in the C terminus was also amplified by PCR and further cloned in similar pCAGGS plasmid. The coding sequences of human Tollip and IFNGR1 were obtained from GenScript (catalog number OHU02397D) and Sino Biological (catalog number HG10338-CF), respectively. These were cloned with an N-terminal Flag tag into the pCAGGS plasmid. Tollip mutants Tollip_1-229_, Tollip_1-180_, Tollip_54-274_, Tollip_54-229_, Tollip_54-180_, Tollip_180-274_, Tollip_180-229_, Tollip_WTHI-AAAA_, Tollip_WYSL-AAAA_, Tollip_WTHI-AAAA/WYSL-AAAA_, Tollip_M240A_,Tollip_F241A_, Tollip_L267A_, and Tollip_L268A_ were cloned in the same manner. Deoxyoligonucleotide sequences of primers used are provided in Supplementary Table 1. The resulting constructs were validated by Sanger DNA sequencing prior to use. Other plasmids used include an NF-κB-responsive firefly luciferase reporter plasmid pGL4.32 [luc2P/NF-κB RE/Hygro] (Promega), an IFN stimulated response element firefly luciferase reporter plasmid pGL4.45 [luc2P/ISRE/Hygro] (Promega), a gamma activated sequence firefly luciferase reporter plasmid (pGL4[luc2P/GAS RE Hygro] (Promega), and a constitutively expressing Renilla luciferase reporter plasmid pRL-TK (Promega).

### DNA transfection and dual luciferase assays

DNA transfection was performed using Lipofectamine 2000 (Invitrogen catalog number 11668500) according to the manufacturer’s instructions. Cells were seeded in 96 well plates. In each well, 0.1 μg of pNF-κB-Luc, pISRE-Luc, or pGAS-Luc, or 0.1 μg of pRL-TK, and 0.3 μg of protein expression plasmid were co-transfected. For the reporter gene assays, at 24 h post-transfection, cells were stimulated with 100 ng LPS for the NF-κB luciferase reporter, 1000 IU/ml of universal IFNα for the ISRE luciferase reporter, 500 IU/ml of human interferon-gamma (IFN-γ) for the gamma activated sequence reporter for 18h and lysates were prepared using Passive lysis buffer (Promega). Supernatants were collected and assayed for firefly and *Renilla* luciferase activities using the Dual luciferase reporter assay system (Promega E1960) and Agilent BioTek Cytation C10 Confocal Imaging Reader. Values for firefly luciferase reporter activities were normalized to the *Renilla* luciferase control, and results were expressed as fold increase versus the untreated, empty vector control. Each transfection was performed in triplicate, and each experiment was repeated at least twice.

### Co-immunoprecipitation assays

HEK293T cells in 6 well plates were transfected with the indicated plasmids by using Lipofectamine 2000. After 48 h incubation, the cells were washed once with PBS, lysed in NP40 lysis buffer (50 mM Tris-HCl pH 8, 280 mM NaCl, 0.5% NP-40, 0.2 mM EDTA, 2 mM EGTA, Glycerol 10%, Protease inhibitors (Sigma-Aldrich catalog number 11836170001)) and centrifuged at 21,130 rcf in an eppendorf 5424 R microcentrifuge for 10 min at 4°C. The resulting supernatants were considered whole cell extracts. Whole cell extracts were incubated with anti-HA magnetic beads (Pierce™ Anti-HA Magnetic Beads catalog number 88836) for 45 minutes at 4°C. A portion of supernatant was saved as input material. The beads were washed five times with NP40 lysis buffer. The IP complex was resuspended in 1X SDS-PAGE loading buffer and heated to 95^0^C for 5 minutes.

### Immunoblotting

Proteins were separated by SDS-PAGE gels (Invitrogen™ Bolt™ 4 to 12%, Bis-Tris catalog number NW04120BOX) and subsequently transferred onto a PVDF membrane (Sigma-Aldrich catalog number 03010040001). The membrane was blocked with 5% blocking buffer (Bio-Rad catalog number 1706404) for 1 h at room temperature followed by overnight incubation with the indicated antibodies, and incubated with Western Chemiluminescent HRP substrate (Perkin Elmer catalog number NEL104001EA). Imaging was performed with a ChemiDoc™ MP Imaging System (Bio-Rad). Detailed information for all antibodies is provided in Supplementary Table 3.

### Deglycosidase treatments

In step 1, 5 µg of HEK293T-expressing Nsp14 protein lysate was mixed with 1 µL of 10X Glycoprotein Denature Buffer and sufficient H_2_O to achieve a total volume of 10 µL. The mixture was boiled at 100°C for 10 minutes to denature the glycoproteins. In step 2, for Endo H treatment: in a new tube, 2 µL of 10X GlycoBuffer 3, 2 µL of Endo H (New England Biolabs catalog number P0702L), were combined with H_2_O to reach a total volume of 10 µL. For PNGase F Treatment: in another tube, 2 µL of 10X GlycoBuffer 2, 2 µL of 10% NP-40, 2 µL of PNGase F (New England Biolabs catalog number P0704S), were combined with H_2_O to achieve a total volume of 10 µL. The denatured sample from Step 1 was added to this mix and incubated at 37 °C for 1 hour. Treated and mock-treated samples were analyzed by western blot.

### RNA isolation and quantitative reverse transcription PCR (RT-qPCR)

RNA was isolated using the DirectZol RNA kit (Zymo Research R2052) according to the manufacturer’s instructions. RNA was reverse transcribed using the SuperScript™ IV VILO™ Master Mix with ezDNase™ Enzyme (catalog number 11766050) based on the manufacturer’s protocol. The deoxyoligonucleotide PCR primer sequences are as listed in Supplementary Table 2. Real time qPCR was conducted using PerfeCTa® SYBR® Green FastMix® (catalog number 101414-272). Quantitative PCR reactions were performed with a Bio Rad CFX Opus 96 qPCR machine under the following condition: 95°C for 10 m, 40 cycles of 95°C for 15 sec, 60°C for 1 min. Relative gene expression for target gene mRNAs were normalized to β-actin mRNA, and the fold difference in gene expression was calculated using the threshold cycle (ΔΔCT) method.

### Immunofluorescence assay (IFA)

Huh7 cells were grown on coverslips for 16 h. Cells were transfected with 0.5 μg of plasmid DNA for 24 h. Cells were fixed with 4% paraformaldehyde in phosphate buffered saline (PBS) supplemented with 1 mM CaCl_2_ and 1 mM MgCl_2_ (PBS-CM) for 15-30 min at room temperature (RT). Fixed cells were washed twice with PBS-CM and permeabilized for 10 min with 0.1% Triton X-100 in PBS-CM at RT. After blocking with 4% goat serum (MP Biomedicals, 0219135680) in PBS supplemented with 0.5% BSA and 0.15% Glycine (PBG) for 1h at RT, cells were incubated with the primary antibodies in blocking buffer for 1h at RT, followed by three washes with PBS for 5 min each and incubation with the secondary antibodies for 1h at RT. The cells were washed three times with PBS and the coverslips were mounted with ProLong™ Glass Antifade Mountant with NucBlue™ Stain (Thermo Fisher Scientific P36981). Stained cells were examined under a Zeiss LSM980 confocal laser scanning microscope. For quantification, images were analyzed with Imaris software (imaris.oxinst.com/), counting at least 50 cells per sample.

### siRNA gene silencing

A549-ACE2 cells were seeded in 6-well plates at a density of 100,000 cells/well. The next day cells were transfected with 50 nM of Tollip SMARTpool (Dharmacon catalog number L-016930-00-0005), or NON-TARGETplus Non-targeting Control siRNAs siRNA (Dharmacon, catalog number D-001810-01-05) using Lipofectamine RNAiMAX (Invitrogen, 13778-150). Twenty-four hours post-siRNA transfection, cells were infected with SARS-CoV-2.

## Acknowledgments

This work was supported by NIH grants AI161104, AI164080 and AI120943. We thank Emma Komers for expert technical support. We thank Anastasija Čupić and Adolfo Garcίa-Sastre for providing virus and assisting in generating virus stocks.

---

**Supplementary Figure 1.**
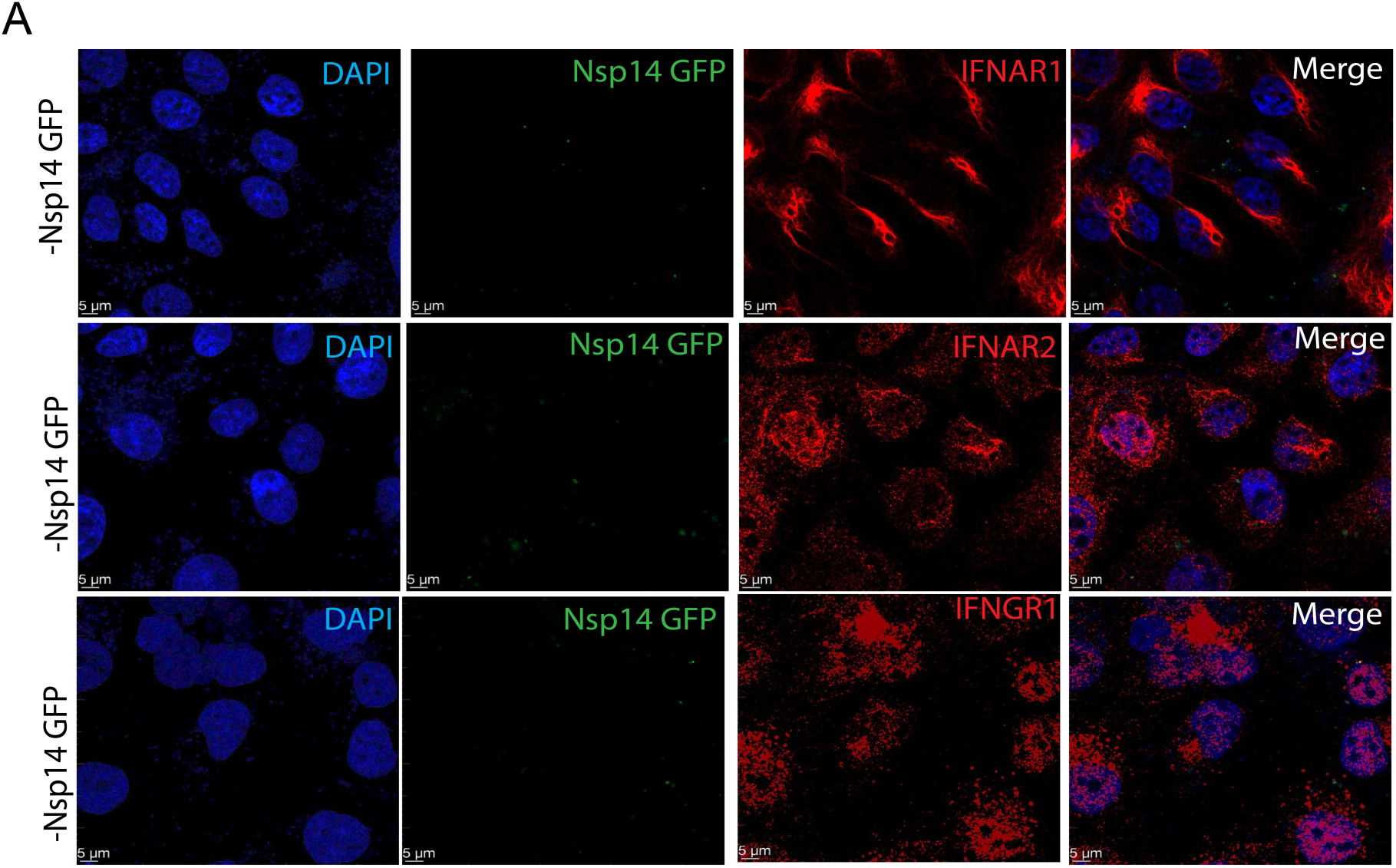
Empty vector control transfections to assess impact on IFNAR1 and IFNGR1 levels. Confocal laser scanning microscopy image of indicated interferon receptor expression level in the presence of empty vector transfected Huh7 cells. Blue, DAPI (nuclei); Green, absence of signal due to absence of Nsp14-GFP; Red, IFNAR1, IFNAR2 or IFNGR1.

**Supplementary Figure 2.**
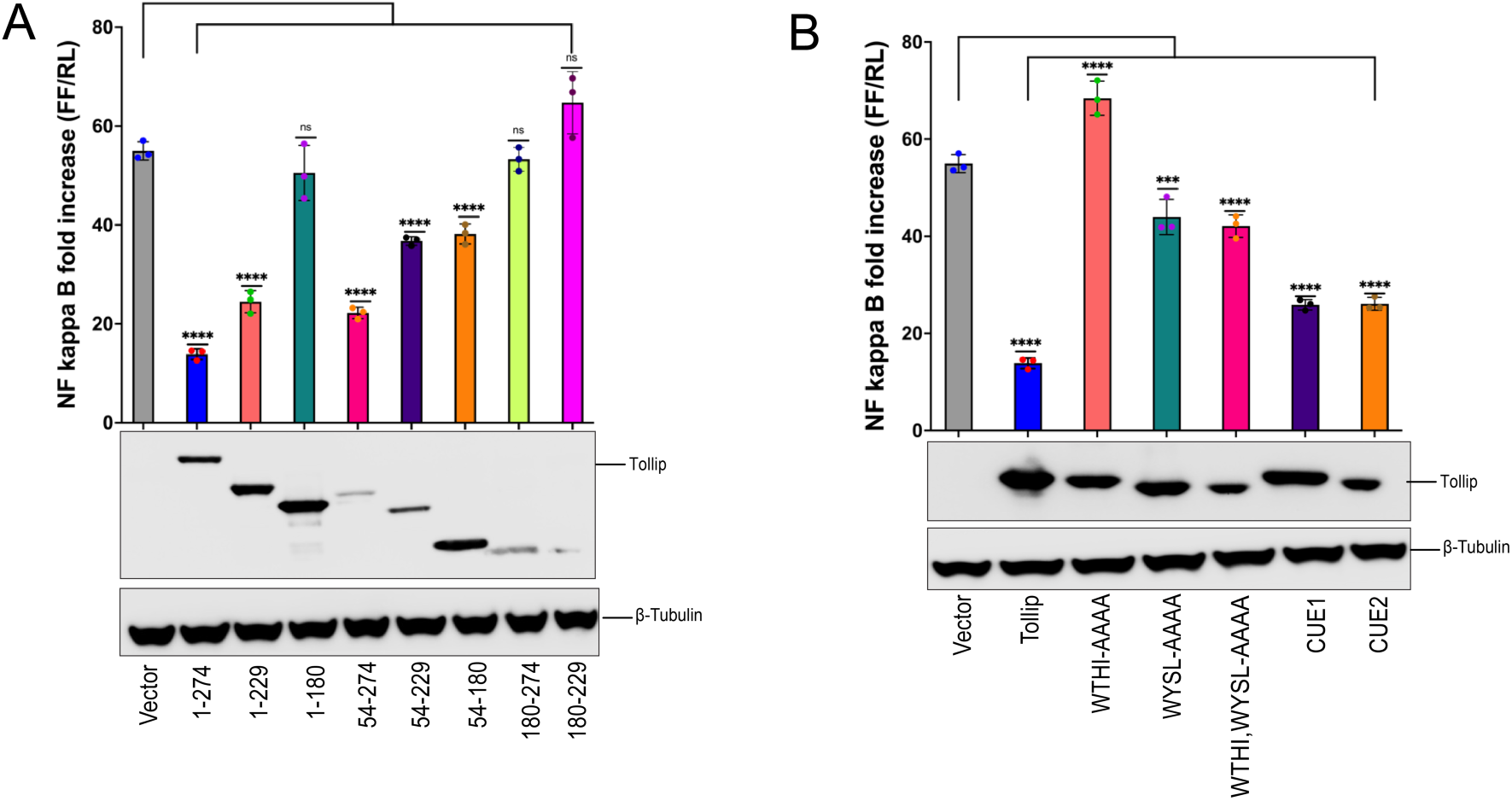
Tollip counteracts LPS-mediated NF-κB activation in the absence of Nsp14. A. Domain mapping studies. HEK293-hTLR4-MD2-CD14 cells were transfected with an NF-κB-firefly luciferase reporter, a constitutively expressing *Renilla* luciferase reporter and the indicated expression plasmids. At 24 h post-transfection, cells were treated with LPS for 18 h and used for dual luciferase reporter assay. Firefly luciferase activity was normalized to *Renilla* luciferase activity. Data are reported as fold increase relative to a mock-treated, empty vector control. Error bars represent mean ± SD (n = 3). One way ANOVA was used to determine statistical significance (P < 0.0001 = ∗∗∗∗, not significant = ns). Cell lysates were analyzed by western blot. B. Tollip point mutant studies. An NF-κB-firefly luciferase reporter assay was performed as in A but with the indicated expression plasmids. (P < 0.001 = ∗∗∗, P < 0.0001 = ∗∗∗∗). Cell lysates were analyzed by western blot.

**Supplementary Table 1.**
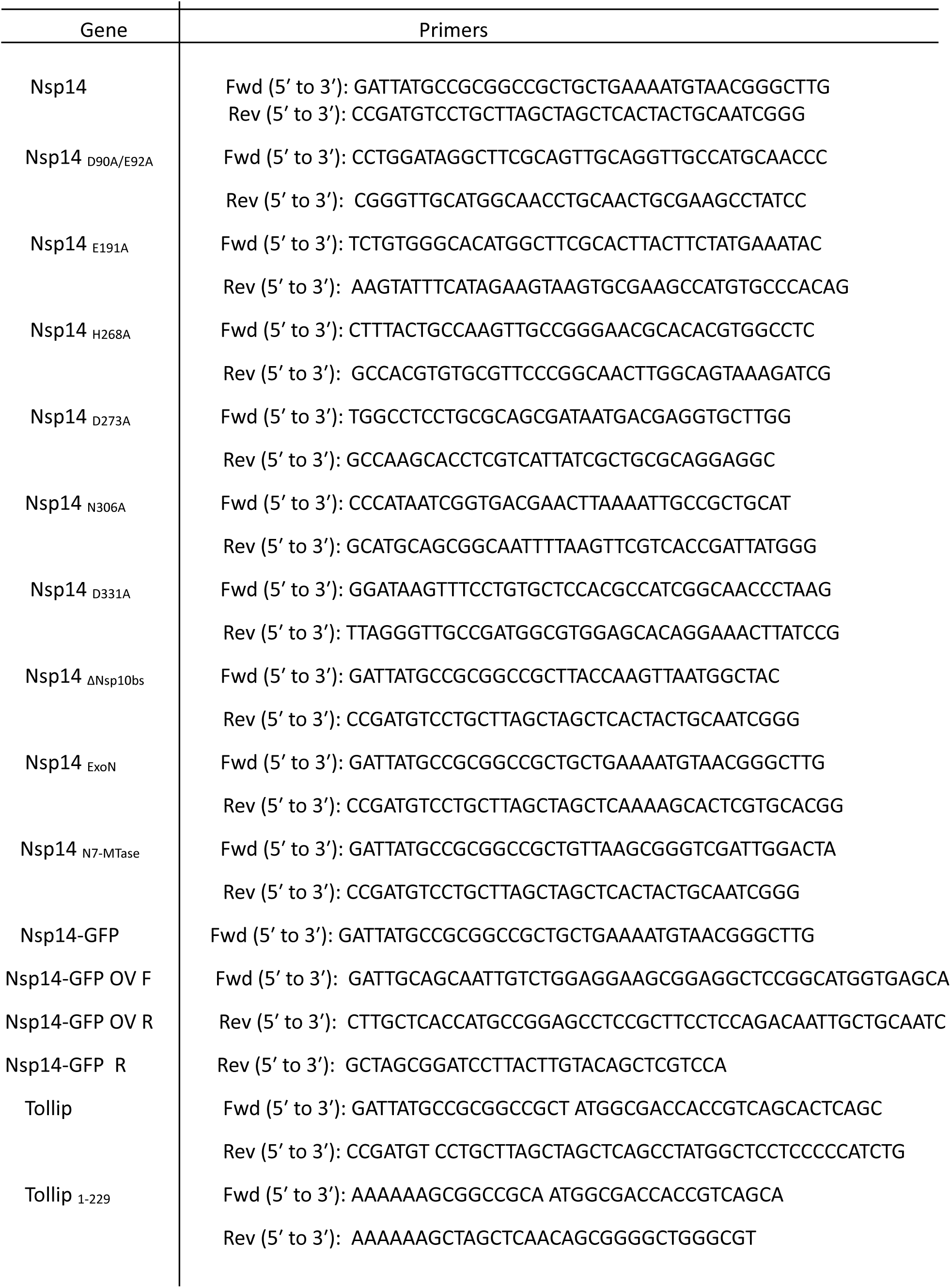
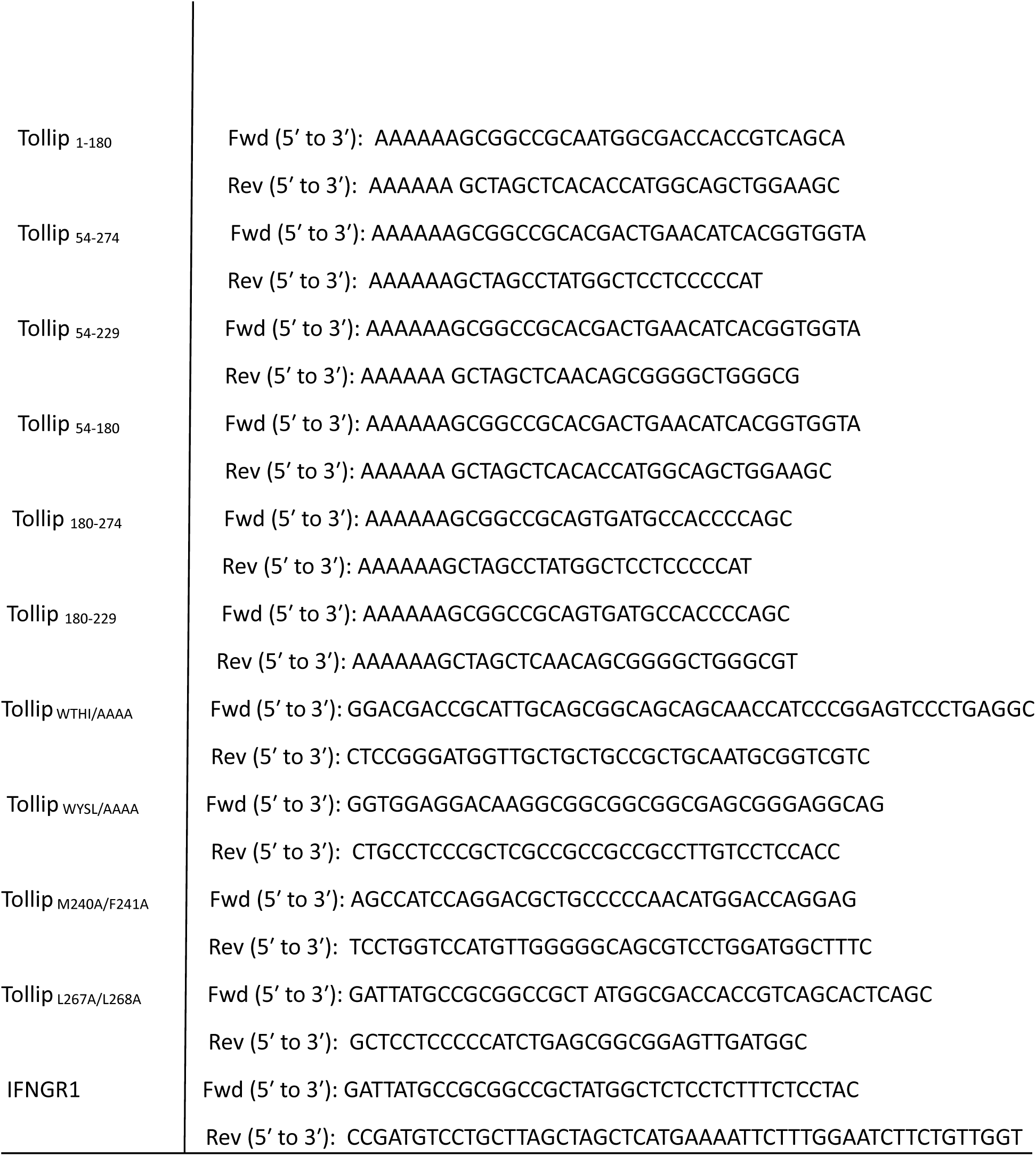
Sequences of primers in this study.

**Supplementary Table 2.**
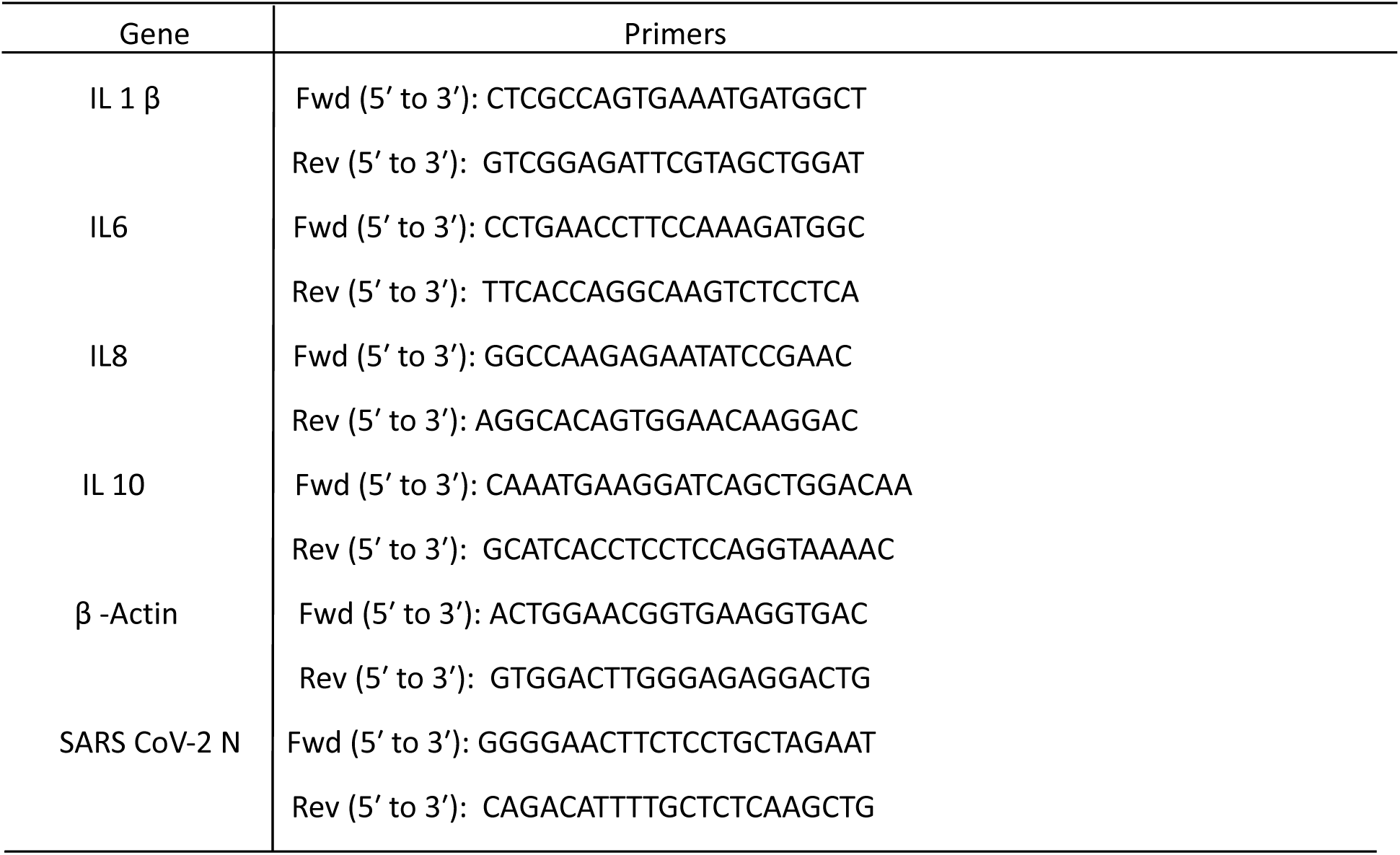
Sequences of primers for qPCR analysis of this study.

**Supplementary Table 3.** Antibodies used in this study.

| Protein | Antibodies |
| --- | --- |
| Tollip | Abcam, (ab187198), anti Tollip Rabbit polyclonal |
| p38 | Cell Signaling Technology (9212S), anti p38 Rabbit monoclonal |
| Phospho-p38 | Cell Signaling Technology (4511S) anti Phospho-p38 Rabbit monoclonal |
| ERK | Cell Signaling Technology (9102S), anti ERK Rabbit polyclonal |
| Phospho-ERK | Cell Signaling Technology (9101S), anti Phospho-ERK Rabbit polyclonal |
| JNK | Cell Signaling Technology (9252S), Anti JNK Rabbit polyclonal |
| Phospho-JNK | Cell Signaling Technology (9151S), anti Phospho-JNK Rabbit polyclonal |
| IFNAR1 | Abcam (ab124764), anti IFNAR1 Rabbit monoclonal (Western blot) |
| IFNAR1 | Abclonal, (A18594), anti IFNAR1 Rabbit polyclonal (IFA) |
| IFNAR2 | Abcam (ab56070), anti IFNAR2 Rabbit polyclonal |
| IFNGR1 | R & D Systems (MAB6731) anti IFNGR1 Mouse monoclonal |
| STAT1 | BD Biosciences (610186), anti STAT1 Mouse monoclonal |
| Phospho-STAT1 | Cell Signaling Technology (9167S), anti Phospho-STAT1 Rabbit monoclonal |
| GAPDH | Thermo Fisher Scientific (MA5-15738), anti GAPDH Mouse monoclonal |
| $\beta$ Tubulin | Sigma (T8328), anti $\beta$ Tubulin Mouse monoclonal |
| Flag-tag | Sigma-Aldrich (F3165), anti-Flag Mouse monoclonal |
| HA-tag | Sigma-Aldrich (H3663) anti HA Mouse monoclonal |
| Flag-tag | Sigma-Aldrich(F7425), anti Flag Rabbit polyclonal |
| HA-tag | Invitrogen (71-5500), anti HA Rabbit polyclonal |
| Anti-mouse IgG | Cell Signaling Technology (7076), Anti-mouse IgG, HRP-linked Antibody |
| Anti-rabbit IgG | Cell Signaling Technology (7074), Anti-rabbit IgG, HRP-linked Antibody |
| Alexa Fluor™ 647 | Invitrogen (A-21236), Goat anti-Mouse, Secondary Antibody, Alexa Fluor™ 647 |
| Alexa Fluor™ 594 | Invitrogen (A-32740), Goat anti-Rabbit Secondary Antibody, Alexa Fluor™ Plus 594 |

